# A consensus definition for deep layer 6 excitatory neurons in mouse neocortex

**DOI:** 10.1101/2024.11.04.621933

**Authors:** Su-Jeong Kim, Travis A. Babola, Kihwan Lee, Chanel J. Matney, Alina C. Spiegel, Michael H. Liew, Eva M. Schulteis, Austin E. Coye, Mikhail Proskurin, Hyunwook Kang, Julia A. Kim, Maxime Chevée, Kiwoong Lee, Patrick O. Kanold, Loyal A. Goff, Juhyun Kim, Solange P. Brown

**Affiliations:** Solomon H. Snyder Department of Neuroscience, Johns Hopkins University School of Medicine, Baltimore, Maryland, 21205, USA; Department of Psychiatry and Behavioral Sciences, Johns Hopkins University School of Medicine, Baltimore, Maryland, 21205, USA; Kavli Neuroscience Discovery Institute, Johns Hopkins University School of Medicine, Baltimore, Maryland, 21205, USA; McKusick-Nathans Institute for Genetic Medicine, Johns Hopkins University School of Medicine, Baltimore, Maryland, 21205, USA; Emotion, Cognition and Behavior Research Group, Korea Brain Research Institute, Daegu, 41062, Republic of Korea; Department of Biomedical Engineering, Johns Hopkins University School of Medicine, Baltimore, Maryland, 21205, USA

## Abstract

To understand neocortical function, we must first define its cell types. Recent studies indicate that neurons in the deepest cortical layer play roles in mediating thalamocortical interactions and modulating brain state and are implicated in neuropsychiatric disease. However, understanding the functions of deep layer 6 (L6b) neurons has been hampered by the lack of agreed upon definitions for these cell types. We compared commonly used methods for defining L6b neurons, including molecular, transcriptional and morphological approaches as well as transgenic mouse lines, and identified a core population of L6b neurons. This population does not innervate sensory thalamus, unlike layer 6 corticothalamic neurons (L6CThNs) in more superficial layer 6. Rather, single L6b neurons project ipsilaterally between cortical areas. Although L6b neurons undergo early developmental changes, we found that their intrinsic electrophysiological properties were stable after the first postnatal week. Our results provide a consensus definition for L6b neurons, enabling comparisons across studies.

## Introduction

To understand the synaptic organization and function of the neocortex, it is essential to establish consensus definitions for its cell types (Ascoli et al., 2008; Yuste et al., 2020). Excitatory neurons residing in the deepest layer of the cortex, often termed layer 6b (L6b) neurons, have been proposed to play roles in mediating thalamocortical interactions and in modulating brain state but results across studies have not been consistent (Ansorge et al., 2020; Arimatsu et al., 2003; Ben-Simon et al., 2022; Hoerder-Suabedissen et al., 2018; Ledderose et al., 2023; Tiong et al., 2019; Viswanathan et al., 2017; Yoneda et al., 2023; Zolnik et al., 2024; Zolnik et al., 2020). As studies define L6b neurons using a variety of approaches (Ansorge et al., 2020; Ben-Simon et al., 2022; Clancy and Cauller, 1999; Gouwens et al., 2019; Heuer et al., 2003; Hoerder-Suabedissen et al., 2018; Hoerder-Suabedissen et al., 2013; Hoerder-Suabedissen et al., 2009; Ledderose et al., 2023; Marx and Feldmeyer, 2013; Reep, 2000; Reep and Goodwin, 1988; Scala et al., 2021; Tasic et al., 2016; Tasic et al., 2018; Tiong et al., 2019; Vandevelde et al., 1996; Venkatesan et al., 2023; Viswanathan et al., 2017; Yao et al., 2021; Yoneda et al., 2023; Yu et al., 2019; Zolnik et al., 2024; Zolnik et al., 2020), we hypothesized that different populations of neurons are captured by the array of methods used, contributing to conflicting results related to their synaptic organization and function.

Some studies defined L6b neurons by their location relative to the white matter border below the neocortex (Chang et al., 2024; Marx and Feldmeyer, 2013; Marx et al., 2017; Yildirim et al., 2019; Zolnik et al., 2020) or by injections of retrograde neuronal tracers into superficial neocortex (Clancy and Cauller, 1999). Others used molecular markers such as Connective Tissue Growth Factor (CTGF), Complexin 3 (Cplx3) or Neurexophilin 4 (Nxph4) to identify L6b neurons (Chang et al., 2024; Heuer et al., 2003; Hoerder-Suabedissen et al., 2018; Hoerder-Suabedissen et al., 2009; Viswanathan et al., 2017; Zolnik et al., 2024; Zolnik et al., 2020). Yet other studies relied on Cre recombinase (Cre) expression in transgenic mouse lines like the Drd1a-Cre line to investigate L6b neurons (Chang et al., 2024; Hoerder-Suabedissen et al., 2018; Zolnik et al., 2024; Zolnik et al., 2020). The contradictory results across these studies, including, for example, differences over whether L6b neurons project to the thalamus and modulate thalamic function (Hoerder-Suabedissen et al., 2018; Ledderose et al., 2023; Tiong et al., 2019; Viswanathan et al., 2017; Zolnik et al., 2024; Zolnik et al., 2020), raise the possibility that the varied methods for defining L6b neurons delineate different populations of neurons with divergent effects on brain function.

To test this possibility, we directly compared commonly used methods for identifying L6b neurons in adult mice. Several methods, including retrograde labeling from superficial cortical layers and expression of CTGF, Cplx3 and Nphx4, defined similar neuronal populations. Using these methods, we developed a consensus definition, and found that L6b neurons were morphologically similar, multipolar excitatory neurons that maintained consistent electrophysiological properties from juvenile to adult mice. Single L6b neurons interconnected ipsilateral cortical areas but did not project to contralateral cortex nor to thalamic nuclei. In addition, L6b neurons in visual cortex of adult mice received little geniculocortical input, consistent with studies in somatosensory cortex (Zolnik et al., 2020), further indicating that they are not strongly integrated into primary sensory thalamocortical circuits. In contrast, in Drd1a-Cre mice, Cre was expressed in only a small subset of L6b neurons defined using our consensus definition. Conversely, a substantial fraction of retrogradely labeled layer 6 corticothalamic neurons (L6CThNs) with typical L6CThN cell body and dendritic morphology expressed Cre, explaining why studies using Drd1a-Cre mice report both anatomical and functional interactions with thalamus (Ansorge et al., 2020; Hoerder-Suabedissen et al., 2018; Zolnik et al., 2024; Zolnik et al., 2020). Together, our results provide a consensus definition for the dominant excitatory neuron type in deep L6 of adult mice, allowing the field to resolve conflicting findings and enabling more precise interrogations of their function.

## Results

### Neurons retrogradely labeled from superficial cortical layers express Connective Tissue Growth Factor (CTGF) and Complexin 3

Numerous methods have been used to identify L6b neurons including retrograde neuronal tracers, anatomical location relative to the white matter, molecular markers and transgenic mouse lines (Clancy and Cauller, 1999; Heuer et al., 2003; Hoerder-Suabedissen et al., 2018; Hoerder-Suabedissen et al., 2009; Marx and Feldmeyer, 2013; Marx et al., 2017; Tiong et al., 2019; Viswanathan et al., 2017; Yildirim et al., 2019; Zolnik et al., 2024; Zolnik et al., 2020). Injecting a retrograde tracer into the upper layers of cortex identifies a thin band of retrogradely labeled neurons just above the white matter border in rats (Clancy and Cauller, 1999). Consistent with this work, injecting the retrograde tracer, AlexaFluor-conjugated cholera toxin subunit B (CTB), into layer 1 and upper layer 2 (L1/2) of the somatosensory cortex of juvenile mice labels a distinct population of cells deep in L6, abutting the white matter underlying the cortex, in addition to labeling neurons in layers 2/3 (L2/3) and 5 (L5) (Figure 1Ai, Aiv). The cell bodies of the retrogradely labeled deep L6 cells were typically ovoid in shape rather than the pyramidal shape typical of cortical pyramidal neurons. Labeled cells in L6 were almost exclusively located in the bottom 10% of the somatosensory cortex, as measured from the pia to the border between the cortex and the white matter (Figure 1B).

**Figure 1:**
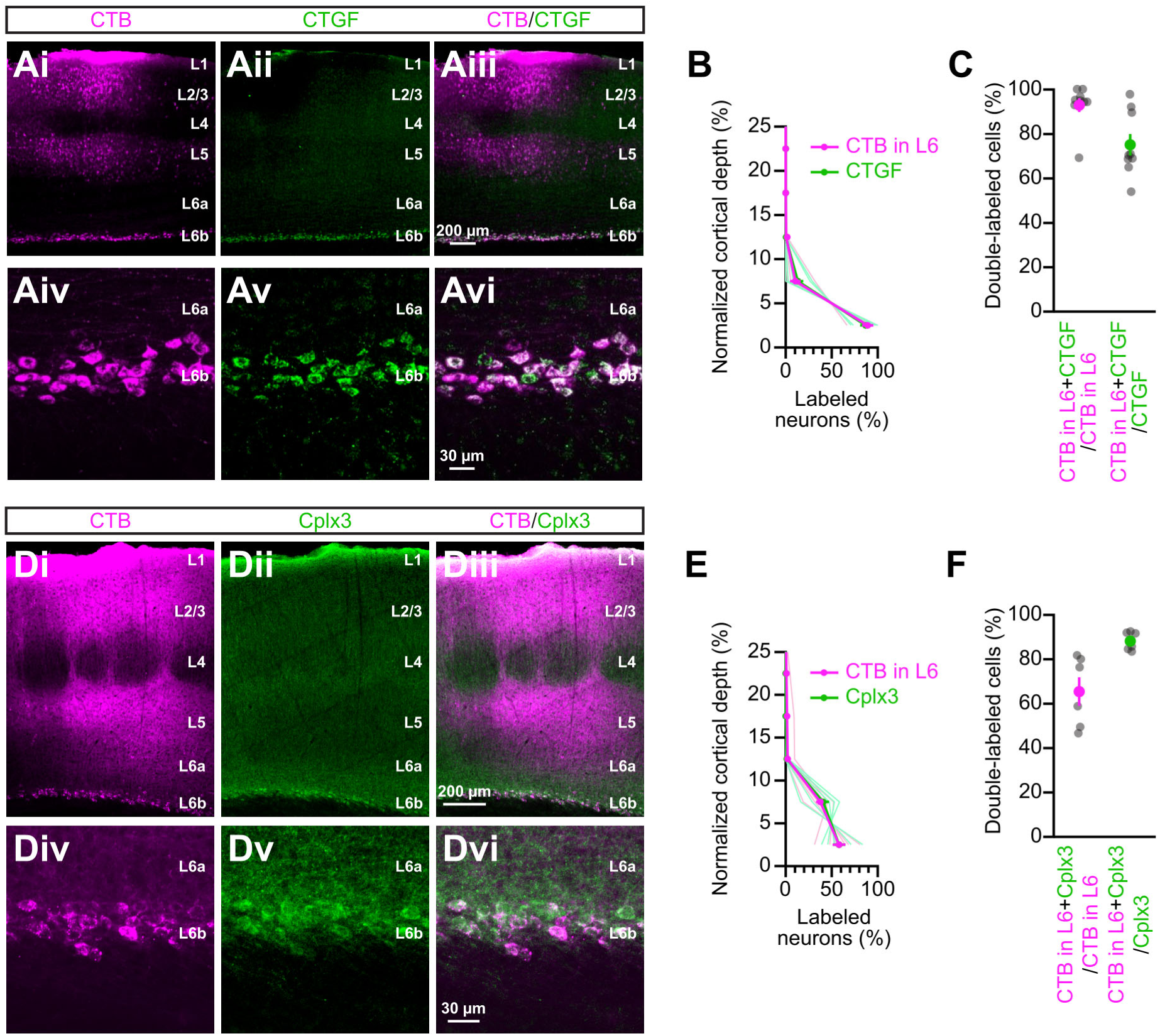
Comparison of deep layer 6 (L6) neuronal populations labeled by local injections of retrograde neuronal tracers and immunostaining for Connective Tissue Growth Factor (CTGF) or Complexin 3 (Cplx3). *A,* Low (Ai-Aiii) and high (Aiv-Avi) magnification images of neurons retrogradely labeled from layers 1 and 2 (L1/2; AlexaFluor 555 Cholera toxin subunit B (CTB); magenta, Ai, Aiv) and immunostained with antibodies to CTGF (green, Aii, Av) in the primary somatosensory cortex (S1) of a postnatal day 26 (P26) mouse. The two images are overlaid (Aiii, Avi) to show double-labeled neurons. *B,* Summary data showing the laminar distribution of retrogradely labeled neurons and CTGF-positive cells in S1 (n = 5 mice, P26-P33; light lines: individual mice; dark line: mean). *C,* Summary data showing the percentage of retrogradely labeled neurons in deep L6 that are CTGF-positive and the percentage of CTGF-positive neurons that are retrogradely labeled in S1 (n = 9 mice, P26-P40). *D,* Low (Di-Diii) and high (Div-Dvi) magnification images of neurons retrogradely labeled from L1/2 (AlexaFluor 647 CTB; magenta, Di,Div) and immunostained with antibodies to Cplx3 (green, Dii, Dv) in S1 of a P30 mouse. The two images are overlaid (Diii,Dvi) to show double-labeled neurons. *E,* Summary data showing the laminar distribution of retrogradely labeled neurons and Cplx3-positive cells in S1 (n = 6 mice, P28-P30; light lines: individual mice; dark lines: mean). *F,* Summary data showing the percentage of retrogradely labeled neurons in deep L6 that are Cplx3-positive and the number of Cplx3-positive neurons that are retrogradely labeled in S1 (n = 6 mice, P28-P30). Scale bars: Ai-iii, Di-ii: 200 μm, Aiv-vi, Div,vi: 30 μm.

Expression of CTGF has also been used to identify a population of neurons in deep L6 across a variety of mammalian species (Heuer et al., 2003; Hoerder-Suabedissen and Molnár, 2013; Hoerder-Suabedissen et al., 2009; Rowell et al., 2010; Sorensen et al., 2015; Viswanathan et al., 2012; Viswanathan et al., 2017; Watakabe et al., 2007). To test the relationship between retrogradely labeled deep L6 neurons and CTGF-positive neurons in the adult mouse, we immunostained neurons retrogradely labeled from L1/2 with antibodies against CTGF (Figure 1Ai-Avi). As with retrogradely labeled neurons in L6, strongly labeled CTGF neurons were primarily located in the bottom 10% of the somatosensory cortex (Figure 1B). Almost all retrogradely labeled neurons in deep L6 were CTGF-positive (Figure 1C). Similarly, the majority of CTGF-positive deep L6 neurons underlying the injection site were also retrogradely labeled by local injections of CTB into L1/2 (Figure 1C). Thus, the two methods identified similar populations of deep L6 neurons.

Another commonly used molecular marker for L6b neurons is Complexin 3 (Cplx3) (Hoerder-Suabedissen and Molnár, 2013; Hoerder-Suabedissen et al., 2009; Viswanathan et al., 2017). We next tested whether immunostaining for Cplx3 identifies a neuronal population similar to the one identified by injections of a retrograde neuronal tracer into L1/2 (Figure 1D). Like the retrogradely labeled neurons, Cplx3-positive neurons were primarily located in the bottom 10% of the somatosensory cortex (Figure 1E). Almost all cells immunopositive for Cplx3 were retrogradely labeled while more than half of the retrogradely labeled deep L6 neurons were Cplx3-positive (Figure 1F). This fraction was significantly smaller than the fraction of retrogradely labeled deep L6 neurons immunopositive for CTGF (p = 0.0024, Mann Whitney U Test), consistent with previous results showing that not all CTGF-positive neurons express Cplx3 (Chang et al., 2024; Viswanathan et al., 2012).

To further compare the neuronal populations expressing *Cplx3* and *Ccn2*, the gene encoding CTGF, we took advantage of recent transcriptomic studies of single cortical cells to assess their expression across cell types (Tasic et al., 2018; Yao et al., 2021). We analyzed gene expression of neurons in three cortical areas: primary visual cortex (V1), anterior lateral motor cortex (ALM) and primary motor cortex (M1) using publicly available transcriptomic data sets (Figure 2). As expected, *Ccn2* was most highly expressed in neurons annotated as L6b neurons in all three cortical areas (Figure 2B). However, in addition to some non-neuronal cell types, excitatory non-projection neurons as well as subpopulations of other cortical excitatory and inhibitory neurons, also expressed *Ccn2*. *Cplx3* was also most highly expressed in neurons annotated as L6b neurons in all three cortical areas (Figure 2C). However, as with *Ccn2*, *Cplx3* expression was also detected in additional cortical neuronal cell types. For example, *Cplx3* was also relatively highly expressed in *Lamp5* inhibitory interneurons. Together, these results indicate that, although high expression of *Ccn2* and *Cplx3* identifies a similar population of neurons in deep L6 as retrograde labeling from L1/2, these two molecular markers are also expressed in additional cortical cell types beyond L6, and thus their expression is not exclusive to deep L6 excitatory neurons.

**Figure 2:**
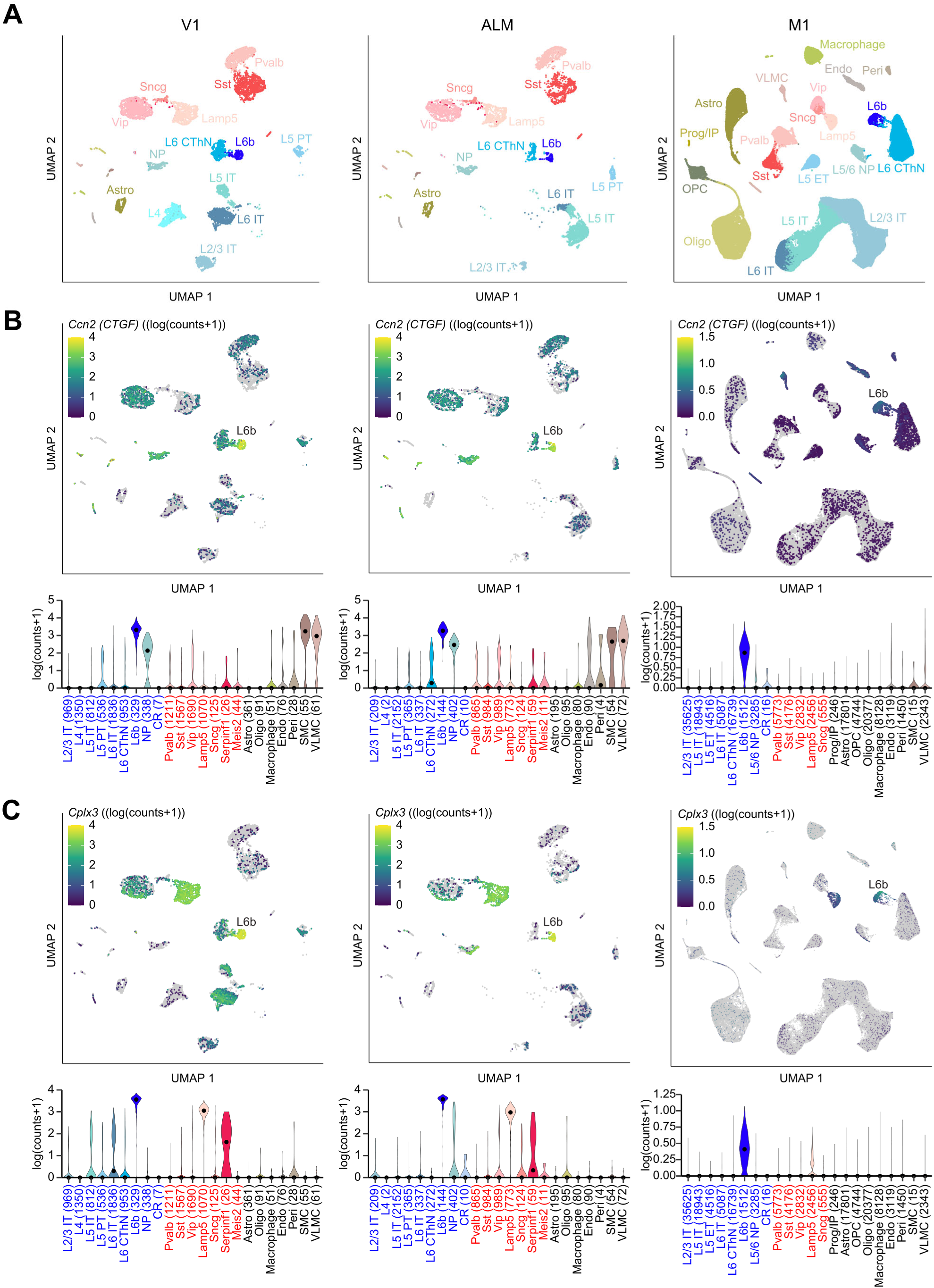
Expression levels of *Ccn2* and *Cplx3* across transcriptionally defined neuronal cell types in primary visual cortex (V1), anterior lateral motor cortex (ALM) and primary motor cortex (M1). *A,* Two-dimensional projection (uniform manifold approximation and projection (UMAP; McInnes et al., 2018) of the transcriptional profiles of adult cortical neurons generated using publicly available data sets from primary visual cortex (V1) (Tasic et al., 2018), anterior lateral motor cortex (ALM) (Tasic et al., 2018) and primary motor cortex (M1) (Yao et al., 2021). UMAPS are colored by cell type annotation with glutamatergic neuron types shown in shades of blue and GABAergic neuron types in shades of red. *B-C*, Gene expression levels for *Ccn2*, the gene encoding CTGF (**B**), and *Complexin 3 (Cplx3,* C) across cell types from the three cortical areas. Gene expression levels are depicted on a log scale with pseudo count of 1, where gray cells have zero counts of the gene. Excitatory neuron subtype annotations: L6b, L6CThN: layer 6 corticothalamic neurons, L2-L6 IT: layer 2-6 intratelencephalic neurons, L5 PT: layer 5 pyramidal tract neurons: NP: near-projecting neurons; Inhibitory neurons subtype annotations: Pvalb, Sst, Lamp5, Scng, and Vip.

### Neurexophilin 4 (Nxph4) expression identifies layer 6b neurons

Neurexophilin 4 (Nxph4) has also previously been used as a molecular marker for deep L6 neurons from early development through adulthood (Hoerder-Suabedissen et al., 2009; Tasic et al., 2018; Wei et al., 2022; Yao et al., 2021), and a Nxph4-CreER mouse line has been used to assess the morphology of L6b neurons (Peng et al., 2021). To compare the population of Cre expressing neurons in this mouse line with the population of deep L6 neurons retrogradely labeled from L1/2 and expressing CTGF and Cplx3, we crossed Nxph4-CreER mice with a tdTomato reporter line, generating mice in which a thin band of neurons, located in the bottom 10% of the neocortex, expressed tdTomato (Figure 3A-D). When we immunostained cortical slices from Nxph4-CreER;tdTomato mice for CTGF, we found that almost all tdTomato-positive neurons were CTGF-positive (Figure 3C-E). However, the efficiency of Cre induction was variable across animals and many CTGF-positive neurons remained tdTomato-negative. Next, using publicly available scRNA-seq data sets (Tasic et al., 2018; Yao et al., 2021), we found that *Nxph4* expression was almost exclusively limited to neurons in the L6b cluster in V1, ALM and M1 (Figure 3F), although there was some expression in pericytes and vascular leptomeningeal cells, consistent with scattered tdTomato expression in vascular-associated cells in some Nxph4-CreER;tdTomato mice. Nonetheless, neuronal *Nxph4* expression was almost exclusively restricted to neurons annotated as L6b neurons in the neocortex. However, neurons outside the neocortex also expressed Cre in this mouse line, including neurons in the cerebellum, a finding consistent with expression in an independently generated Nxph4-βgeo mouse line (Meng et al., 2019). In addition, serotonergic neurons in the brain stem expressed Cre (Figure 3G,H), neurons that may project to the cortex. Thus, although Cre expression is almost exclusively confined to L6b neurons in the cortex, the expression of Cre recombinase in these additional brain areas complicates the use of this mouse line for selective genetic access of L6b neurons through crosses with Cre-dependent mouse lines.

**Figure 3:**
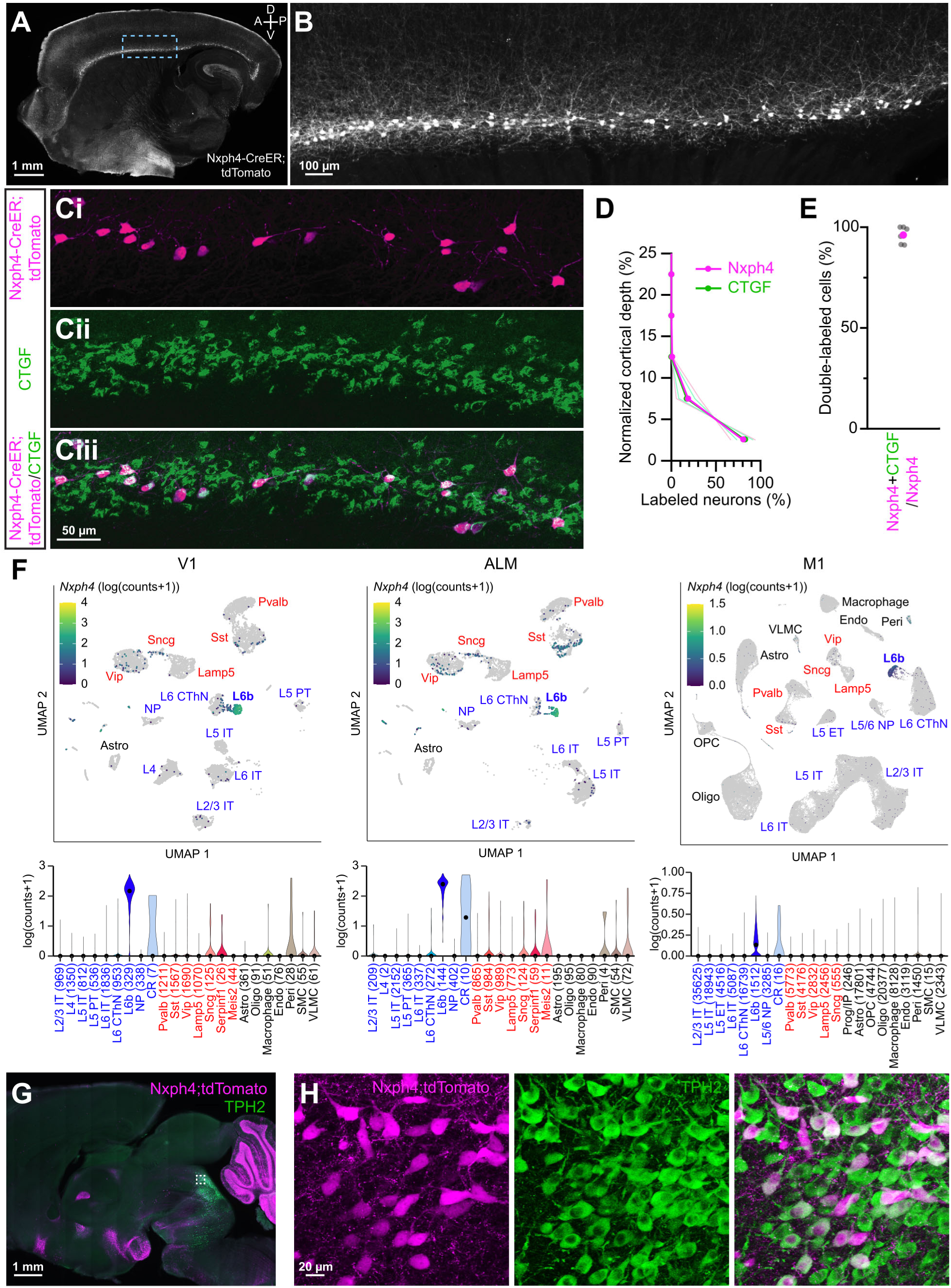
Distribution of Cre expressing neurons in a Neurexophilin 4-CreER (Nxph4-CreER) mouse line identifies layer 6b (L6b) neurons. *A,* Image of a sagittal section from an adult Nxph4-CreER;tdTomato mouse (postnatal day 111, P111). *B,* Higher magnification image of the somatosensory cortex, outlined in blue in *A*, showing tdTomato expression in deep L6 neurons. *C,* Images showing Nxph4-CreER;tdTomato expression (magenta, Ci) and CTGF immunostaining (green, Cii) of deep L6 neurons of somatosensory cortex of a Nxph4-CreER;tdTomato adult mouse (P72). The two images are overlaid (Ciii) to show double-labeled neurons. *D*, Summary data showing the laminar distribution of Nxph4-CreER;tdTomato-positive cells and CTGF-positive cells in S1 (n = 2 mice, P26; light lines: individual mice; dark lines: means). *E*, Summary data showing the percentage of Nxph4-CreER;tdTomato-positive neurons immunopositive for CTGF (n = 6 mice; P32-P147). *F*, Two-dimensional projection (uniform manifold approximation and projection (UMAP; McGuire et al., 1984) of transcriptionally defined neuronal cell types (top) generated using publicly available data sets from primary visual cortex (V1) (Tasic et al., 2018), anterior lateral motor cortex (ALM) (Tasic et al., 2018) and primary motor cortex (M1) (Yao et al., 2021) showing expression levels of *Nxph4* (bottom) in V1, ALM and M1 as in Figure 2. *G*, Low magnification image showing Nxph4-CreER;tdTomato-positive neurons (magenta) in a sagittal section from a P72 mouse immunostained for Tryptophan Hydroxylase 2 (TPH2, green), the rate-limiting enzyme for serotonin synthesis. *H*, Higher magnification image of the region outlined by the white box in *G* showing tdTomato-expression (left), TPH2 immunostaining (middle) and the two overlaid (right). Scale bars: A,G: 1 mm, B: 100 μm, C: 50 μm; H: 20 μm.

### Cre recombinase is expressed in few deep layer 6 neurons in Ntsr1-Cre mice

The Ntsr1-Cre mouse line has been extensively used to genetically access L6CThNs in primary sensory cortical areas (Augustinaite and Kuhn, 2020; Born et al., 2021; Bortone et al., 2014; Chevée et al., 2018; Crandall et al., 2015; Crandall et al., 2017; Dash et al., 2022; Frandolig et al., 2019; Gong et al., 2007; Guo et al., 2017; Kim et al., 2014; Olsen et al., 2012; Spacek et al., 2022; Williamson and Polley, 2019), and several studies have indicated that L6b neurons project to the thalamus (Ansorge et al., 2020; Ben-Simon et al., 2022; Hoerder-Suabedissen et al., 2018; Viswanathan et al., 2017; Zolnik et al., 2024; Zolnik et al., 2020). Therefore, we next tested whether Cre was expressed in deep L6 neurons in this mouse line. When we retrogradely labeled deep L6 neurons from L1/2 in Ntsr1-Cre;tdTomato mice, we found that they formed a band largely below the layer of Ntsr1-Cre;tdTomato-positive neurons (Figure 4A,B). These findings indicate that the tdTomato-negative neurons below the tdTomato-positive L6 neurons in Ntsr1-Cre;tdTomato mice are composed of L6b neurons retrogradely labeled from L1/2, presenting an alternative approach for identifying this cell population. Together, our results indicate that multiple approaches identify similar populations of deep L6 neurons as defined by retrograde labeling from L1/2. These approaches include: 1) CTGF expression; 2) Cplx3 expression; 3) *Nxph4* expression and Cre expression in Nxph4-CreER mice and 4) exclusion of Cre expression in neurons in deep L6 in Ntsr1-Cre mice.

**Figure 4:**
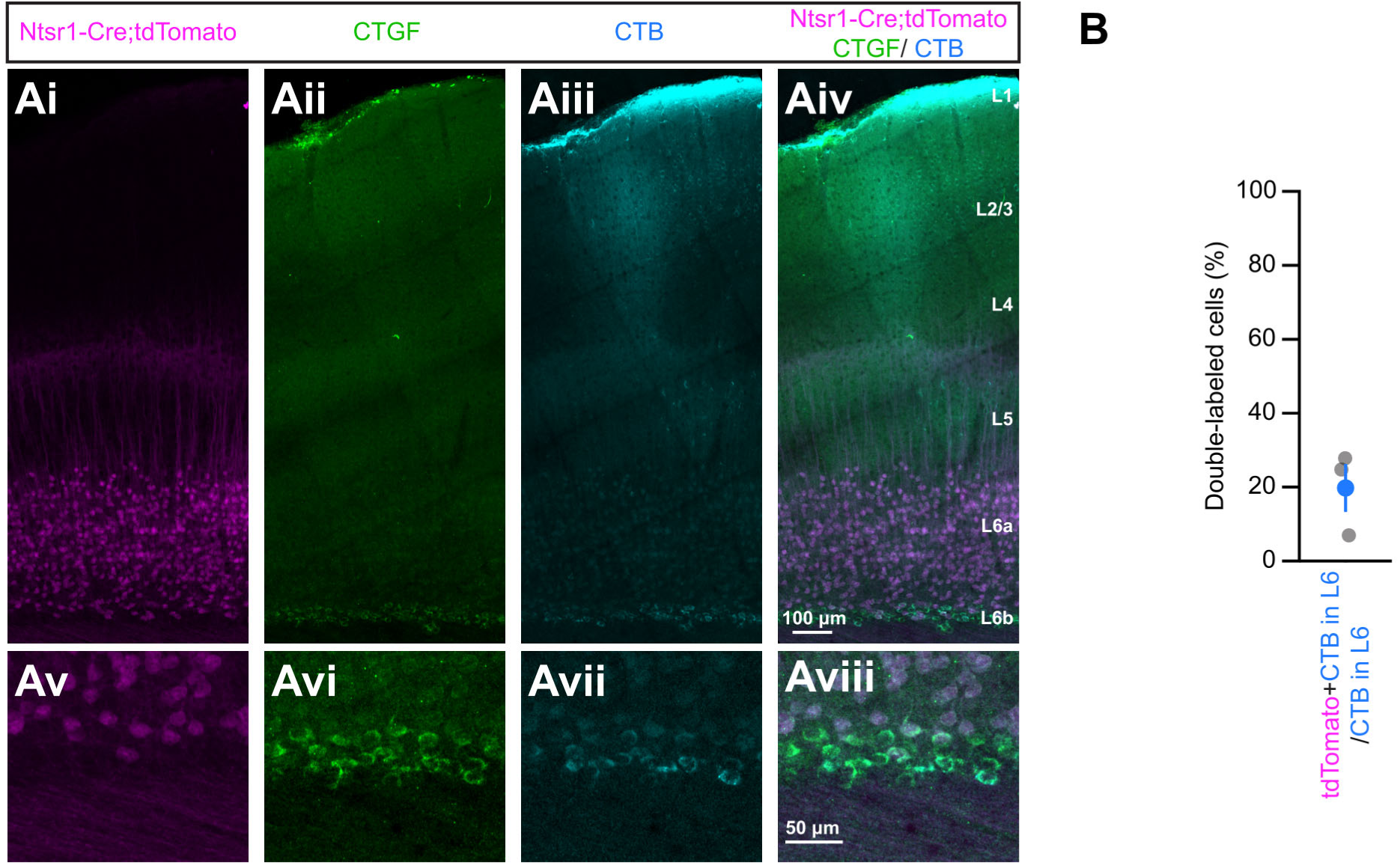
Few layer 6b (L6b) neurons express Cre recombinase in the Ntsr1-Cre mouse line. ***A***, Low (Ai-Aiv) and high (Av-Aviii) magnification images of Ntsr1-Cre;tdTomato-positive neurons (magenta, Ai, Av), CTGF-positive neurons (green, Aii, Avi) and neurons retrogradely labeled from layers 1/2 (L1/2; blue, Aiii, Avii) in S1 of a Ntsr1-Cre;tdTomato mouse (postnatal day 25, P25). The three images are overlaid (Aiv, Aviii). ***B***, Summary data showing the percentage of L1/2 retrogradely labeled neurons that are tdTomato-positive (n = 3 mice, P25-P33) in deep L6 of S1 in Ntsr1-Cre;tdTomato mice. Scale bars: Ai-iiv: 100 μm, Av-viii: 50 μm.

### Layer 6b neurons are spiny, multipolar neurons without apical dendrites

Previous studies indicate that subplate neurons in early development are morphologically heterogeneous (Ghezzi et al., 2021; Hoerder-Suabedissen and Molnár, 2012), including neurons with typical pyramidal cell morphologies as well as cells with ovoid somas and bitufted or multipolar dendritic arbors. Whether L6b neurons in juvenile and adult rodents are similarly heterogeneous is less clear, although studies using different definitions of L6b neurons have identified neurons with apical dendrites, inverted pyramidal neurons and multipolar cells with both smooth and spiny dendrites in the targeted population (Andjelic et al., 2009; Marx and Feldmeyer, 2013; Marx et al., 2017; Peng et al., 2021; Zolnik et al., 2024). We next asked whether the morphology of L6b neurons identified using the consensus definition we developed was heterogeneous. We filled tdTomato-negative neurons in deep L6 of Ntsr1-Cre;tdTomato mice with biocytin and generated three-dimensional reconstructions of the dendritic arbor of L6b neurons in somatosensory and visual cortex. Of the tdTomato-negative neurons below the band of tdTomato-positive L6 neurons we filled with biocytin and immunostained for CTGF (Figure 5A), 15 of 16 neurons were CTGF-positive. We found that these neurons consistently demonstrated multipolar dendritic arbors that were largely confined to L6 (Figure 5A,B). The morphology of these neurons was consistent with the morphology of the population of tdTomato-positive neurons in Nxph4-CreER;tdTomato mice (Figure 3B). We did not identify any neurons with a clear apical dendrite or inverted pyramidal cell morphology. We found that the dendrites of CTGF-expressing deep L6 neurons were studded with dendritic spines, suggesting that they were excitatory cortical neurons (Figure 5C,D). The excitatory nature of these cells was confirmed by immunostaining cortical slices from Gad2-mCherry mice for CTGF. Few double-labeled neurons were identified (S1: 1.7 ± 3.4%; V1: 3.6 ± 7.2%; M1: 1.4 ± 2.8%; n = 4 mice, P70-P81). Consistent with these results and prior studies (Boon et al., 2019; Hoerder-Suabedissen and Molnár, 2013; Hoerder-Suabedissen et al., 2009), we found that deep L6 parvalbumin-positive (PV) and somatostatin-positive (SOM) interneurons, the two most common inhibitory interneurons in the neocortex, did not express CTGF (0 of 106 PV interneurons; 0 of 116 SOM interneurons; n = 9 mice, P52-P223). These results indicate that L6b neurons specified by our consensus definition represent a population of spiny excitatory neurons with multipolar dendritic arbors.

**Figure 5:**
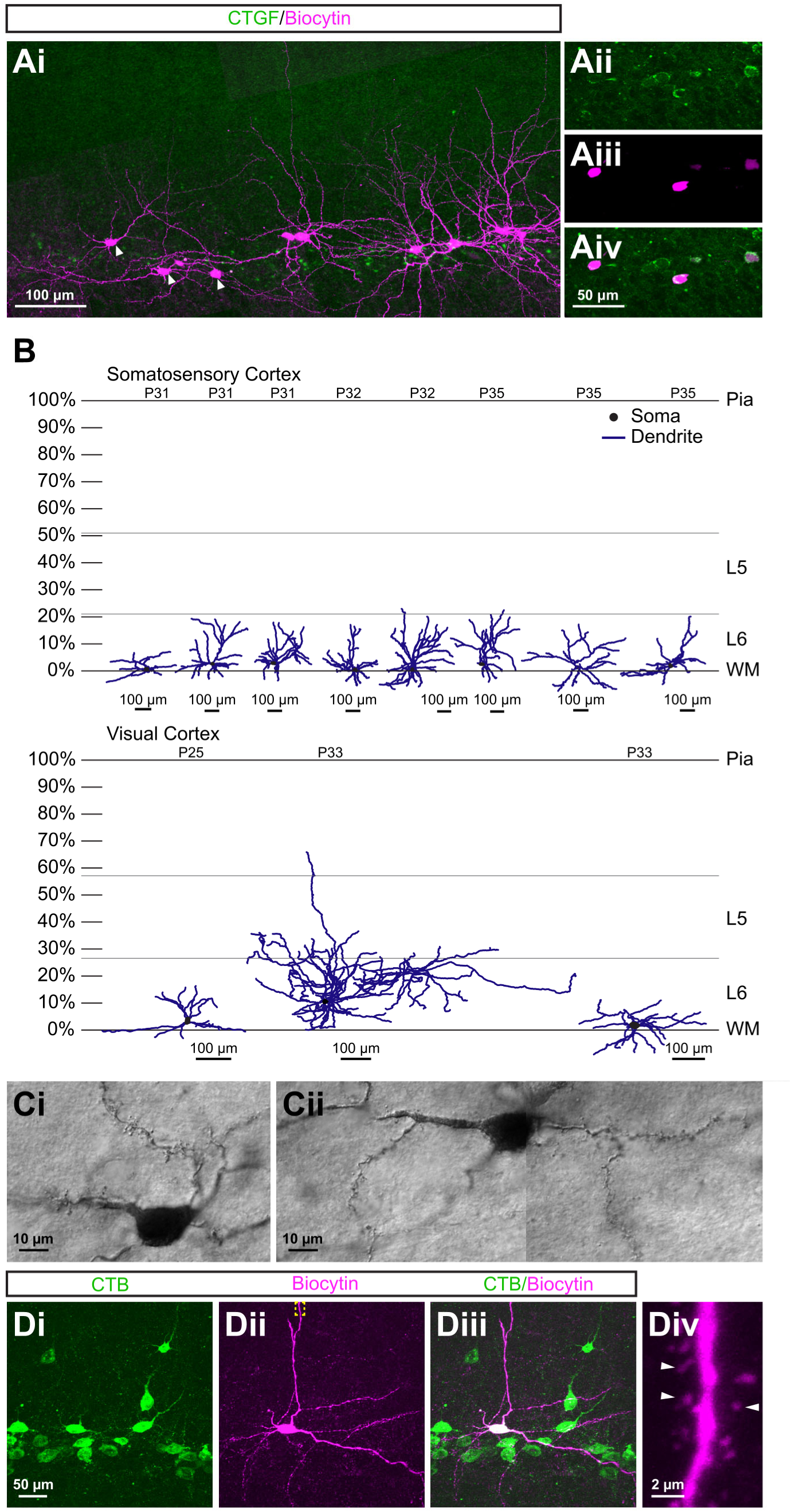
Layer 6b (L6b) neurons are multipolar neurons with spiny dendrites confined to the infragranular layers. ***A,*** Image of deep L6 neurons filled with biocytin (magenta) and immunopositive for CTGF (green) in the somatosensory cortex of a postnatal day 18 (P18) mouse. Higher magnification images of the three cells indicated by arrowheads in (Ai) show CTGF immunostaining (Aii) and biocytin labeling (Aiii). The images are overlaid to show co-localization (Aiv). ***B,*** Three-dimensional reconstructions of biocytin-filled L6b neurons in visual and somatosensory cortex (n = 6 mice; P25-P35; dendrites: blue, somas: black). The vertical extent of the cortex was normalized independently for each reconstruction. ***C,*** Images of the spiny dendrites of a L6b neuron in motor cortex retrogradely labeled from layers 1 and 2 (L1/2) and filled with biocytin (P26 mouse). ***D,*** Images (Di-Diii) showing a biocytin-filled L6b neuron in visual cortex retrogradely labeled from L1/2 in an adult mouse (P46). A higher magnification view of the dendrite outlined by the yellow box (Div). Arrowheads indicate dendritic spines. Scale bars: Ai: 100 μm, Aii-Aiv: 50 μm, B: 100 μm, C: 10 μm, Di-Diii: 50 μm, Div: 2 μm.

### In Drd1a-Cre mice, Cre recombinase is expressed in few layer 6b neurons retrogradely labeled from layers 1 and 2

Drd1a-Cre mice have recently been used to genetically access deep L6 neurons and determine their synaptic organization and contributions to cortical function (Ansorge et al., 2020; Hoerder-Suabedissen et al., 2018; Zolnik et al., 2024; Zolnik et al., 2020). However, the relationship between Cre-expressing neurons in Drd1a-Cre mice and the population of L6b neurons identified using our consensus definition across cortical areas has not been fully resolved. To better understand this relationship, we crossed Drd1a-Cre mice with a Cre dependent tdTomato reporter line and injected one retrograde tracer in L1/2 to retrogradely label L6b neurons and a second tracer in the posterior medial nucleus of the thalamus (POm) to retrogradely label L6CThNs in lower L6 of the somatosensory cortex (Figure 6A) (Bourassa et al., 1995; Chevée et al., 2018; Frandolig et al., 2019; Killackey and Sherman, 2003). We found that Cre-expressing tdTomato-positive neurons were distributed more broadly across L6 than retrogradely labeled L6b neurons (Figure 6B,C). Indeed, the distribution of tdTomato-positive neurons more closely approximated the distribution of retrogradely labeled L6CThNs than L6b neurons (Figure 6C). Of the tdTomato-positive cells, few were retrogradely labeled from L1/2 (6.0 ± 1.9%, n = 4 mice, P26-P32). In contrast, approximately a quarter of the tdTomato-positive cells were L6CThNs retrogradely labeled from PO (25.1 ± 7.3%, n = 4 mice, P26-P32). Injections into contralateral S1 did not result in any double-labeled tdTomato-positive neurons (n = 4 mice, P26-P32). In addition, neither the population of retrogradely labeled L6b neurons nor the L6CThN population were fully captured by Cre expression: only 17.2 ± 3.2% of neurons retrogradely labeled from L1/2 were tdTomato-positive (n = 4 mice, P26-P32). Similarly, only 27.6 ± 5.5% of L6CThNs retrogradely labeled from PO were tdTomato-positive (n = 4 mice, P26-P32). Thus, tdTomato-positive neurons in Drd1a-Cre;tdTomato mice represent a heterogeneous population of cells composed of a small subset of L6b neurons, a larger fraction of POm-projecting L6CThNs and additional cell types.

**Figure 6:**
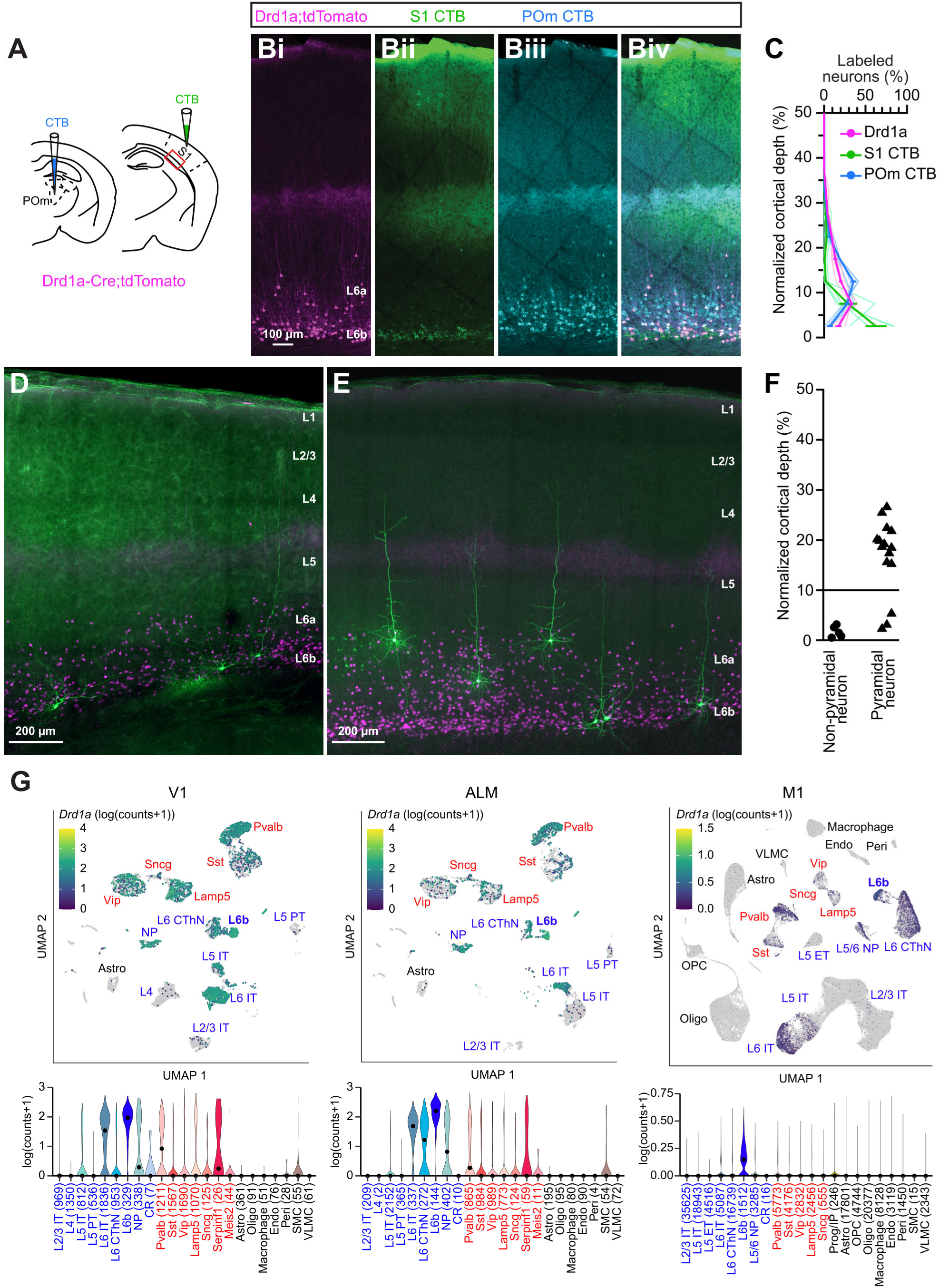
Cre recombinase is expressed in subsets of multiple cell types in a Drd1a-Cre mouse line. ***A,*** Schematic showing the experimental configuration. ***B,*** Image of the somatosensory cortex (S1) of a postnatal day 26 (P26) Drd1a-Cre;tdTomato mouse showing tdTomato-positive cells (Bi), layer 6b (L6b) neurons retrogradely labeled from layers 1 and 2 (L1/2; Bii, green), and corticothalamic neurons retrogradely labeled from the posterior medial nucleus of the thalamus (POm) (Biii, cyan). The three images are overlaid for comparison (Biv). ***C,*** Summary plot showing the laminar distributions of tdTomato-positive neurons, neurons retrogradely labeled from L1/2 and neurons retrogradely labeled from POm in S1 of Drd1a-Cre;tdTomato mice (n = 4 mice; P26-P32; light lines: individual mice; dark lines: means). ***D****,**E**,* Images of tdTomato-positive neurons (magenta) filled with biocytin (green) in S1 of Drd1a-Cre;tdTomato mice (D: P32, E: P25). ***F***, Summary data showing the laminar distribution and morphology of filled tdTomato-positive neurons in somatosensory cortex of Drd1a-Cre;tdTomato mice (n = 20 cells from 6 mice; P23-32). ***G***, Two-dimensional projection (uniform manifold approximation and projection (UMAP; McGuire et al., 1984) of the transcriptional profiles of cortical neurons and expression levels for *Drd1a* in V1, ALM and M1 generated as in Figure 2. Scale bars: B: 100 μm, D,E: 200 μm.

To further confirm the identity of the tdTomato-positive neurons in Drd1a-Cre;tdTomato mice, we next examined their dendritic morphology. We filled individual tdTomato-positive neurons with biocytin in cortical slices from Drd1a-Cre;tdTomato mice. We found that some tdTomato-positive neurons exhibited a multipolar dendritic morphology like L6b neurons identified using our consensus definition (Figure 3A,B), but many had a clear apical dendrite extending into at least L5a (Figure 6D,E). Those tdTomato-positive cells with a multipolar morphology similar to L6b neurons as identified with our consensus definition were generally located closer to the white matter than the tdTomato-positive cells with an apical dendrite (Figure 6F). Of the tdTomato-positive neurons we filled in more superficial L6, all exhibited apical dendrites extending to L5a or more superficially (Figure 6F; n = 12 cells from 4 mice, P23-P32).

Transcriptional analysis of *Drd1a* expression was consistent with these results (Figure 6G). *Drd1a* was expressed not only in L6b neurons but was also highly expressed in neurons annotated as L6CThNs, L6 intratelencephalic (IT) neurons and L5/6 nonprojecting (NP) excitatory neurons as well as in some inhibitory neuron types. Together, these results indicate that Cre is expressed in a heterogeneous population of neurons in Drd1a-Cre mice, composed of L6 pyramidal cell types, including a subset of L6CThNs, as well as a small subpopulation of L6b neurons and some inhibitory interneurons.

### L6b neurons do not project to the thalamus

Whether L6b neurons project to the thalamus has been debated, with important implications for their function. Several recent studies suggest that L6b neurons project to a subset of thalamic nuclei and influence thalamic responses (Ansorge et al., 2020; Ben-Simon et al., 2022; Hoerder-Suabedissen et al., 2018; Viswanathan et al., 2017; Zolnik et al., 2024; Zolnik et al., 2020), while other studies indicate they do not project to the thalamus (Arimatsu et al., 2003). To test whether the population of deep L6 neurons identified using our consensus definition projects to the thalamus, we re-analyzed neurons retrogradely labeled from L1/2 and from POm in sections from Drd1a-Cre;tdTomato mice (Figure 6B). We did not find any neurons double-labeled with both tracers, confirming that L6b neurons retrogradely labeled from L1/2 and L6CThNs retrogradely labeled from POm form distinct populations in the somatosensory cortex. Next, we retrogradely labeled L6CThNs in somatosensory cortex with retrograde tracer injections in POm and immunostained for CTGF (Figure 7A). We found that CTGF-positive neurons were stratified in deep L6, below the layer of POm-projecting L6CThNs (Figure 7B), consistent with our previous work (Frandolig et al., 2019). Very few retrogradely labeled POm-projecting neurons were double-labeled with CTGF (Figure 7C, n= 5 mice, P64-72). To further confirm this point, we retrogradely labeled POm-projecting L6CThNs in Nxph4-CreER;tdTomato mice. Only about 1% of tdTomato-positive neurons were labeled with retrograde tracer (0% and 1.96%, n = 2 mice, aged P72), again confirming that L6b neurons do not project to the thalamus. Similarly, in primary visual cortex, we found that deep L6CThNs projecting to the lateral posterior nucleus (LP), a higher order thalamic nucleus in the visual system targeted by deep L6CThNs (Bourassa and Deschênes, 1995), represent a distinct population from neurons expressing CTGF (Figure 7D,E) with few neurons double-labeled (Figure 7F, n = 4 mice, P24-P27).

**Figure 7:**
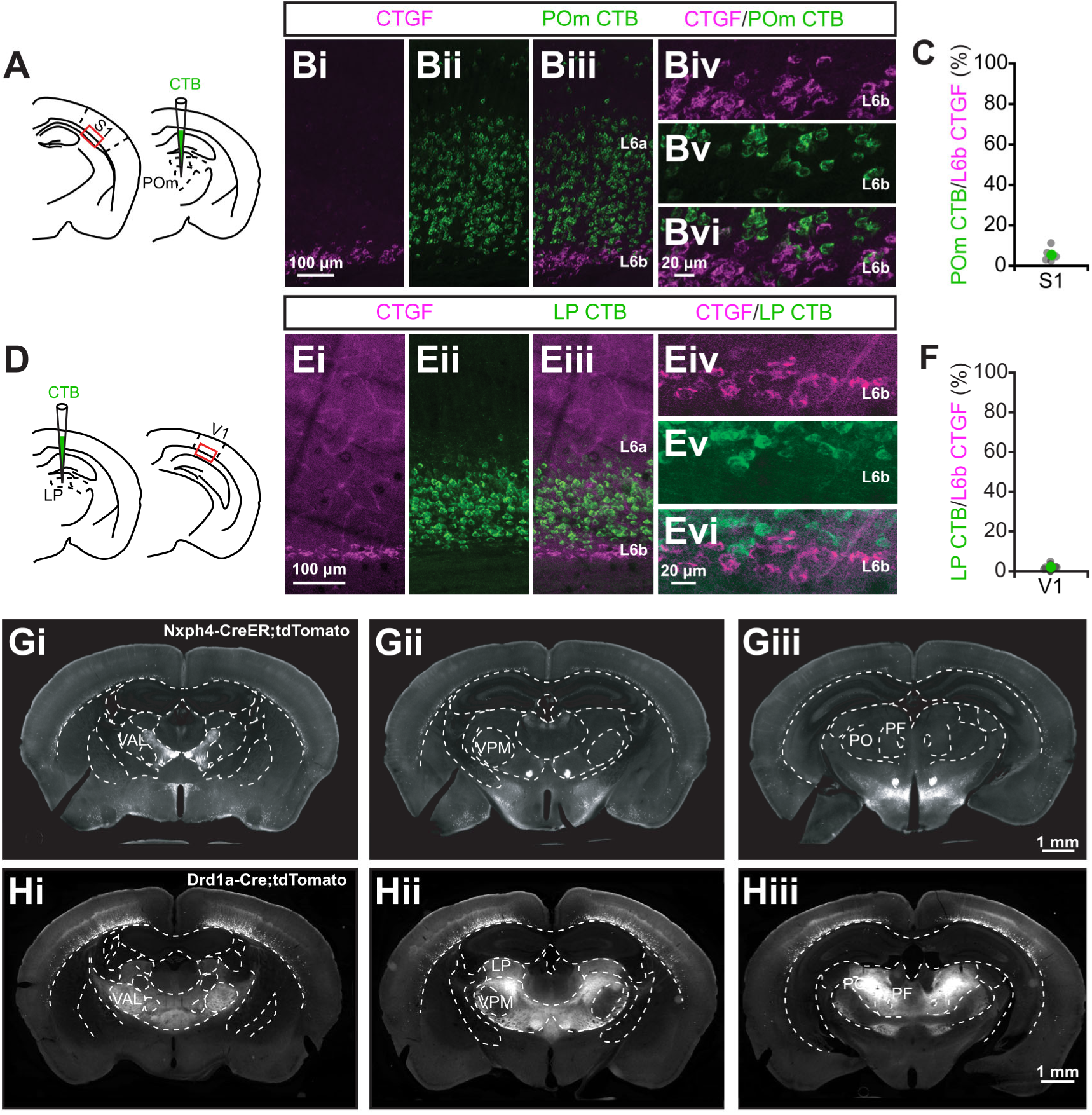
Layer 6b neurons in sensory cortex do not project sensory thalamus. ***A,*** Schematic showing the experimental configuration. ***B,*** Image of the somatosensory cortex immunostained for CTGF (Bi, magenta) and corticothalamic neurons retrogradely labeled from the posterior medial nucleus of the thalamus (POm, green, Bii) from a postnatal day 72 (P72) mouse. The two images are overlaid (Biii). The insets (Biv-Bvi) show deep L6 neurons at higher magnification. ***C,*** Summary data showing the percentage of CTGF-positive neurons retrogradely labeled from POm (n = 5 mice; P64-P72). ***D,*** Schematic showing the experimental configuration. ***E,*** Image of the visual cortex immunostained for CTGF (Ei, magenta) and corticothalamic neurons retrogradely labeled from the lateral posterior nucleus of the thalamus (LP, green, Eii) from a P27 mouse. The two images are overlaid (Eiii). The insets (Eiv-Evi) show deep L6 at higher magnification. ***F,*** Summary data showing the percentage of CTGF-positive neurons retrogradely labeled from LP (n = 4 mice, P24-P27). ***G,*** Three images of coronal sections of a Nxph4-CreER;tdTomato mouse (postnatal day 48, P48) showing a lack of axonal projections into visual or somatosensory thalamus. ***H,*** Three images of coronal sections of a P27 Drd1a-Cre;tdTomato mouse showing axonal projections into higher order visual (LP) and somatosensory (PO) thalamus. Scale bars: Bi-iii,Ei-iii: 100 μm; Biv-vi,Eiv-vi: 20 μm; G,H: 1 mm.

To further confirm these findings, we next crossed Nxph4-CreER mice with a Cre-dependent tdTomato reporter line and examined the thalamus for tdTomato-positive axonal fibers. Although numerous neurons in deep L6 expressed tdTomato, and tdTomato fibers were present in the cortex, thalamic nuclei, including the ventral posterior nucleus (VPM), POm, the lateral geniculate nucleus (LGN), and LP were largely devoid of axonal fibers (Figure 7G). In contrast, tdTomato-positive axonal fibers were observed in thalamic nuclei in Drd1a-Cre;tdTomato mice (Figure 7H), consistent with data showing that a subset of tdTomato-positive neurons in these mice are L6CThNs (Figure 6) and previous reports showing thalamic projections in Drd1a-Cre mice (Ansorge et al., 2020; Ben-Simon et al., 2022; Hoerder-Suabedissen et al., 2018; Viswanathan et al., 2017; Zolnik et al., 2024; Zolnik et al., 2020). Together, these data indicate that L6b neurons defined by retrograde labeling from L1/2 as well as CTGF and Nxph4 expression are not corticothalamic neurons.

### The axons of individual L6b neurons project between cortical areas as well as locally

Previous studies suggest that neurons deep in L6 are retrogradely labeled from tracer injections in neighboring cortical areas (Arimatsu et al., 2003; Clancy and Cauller, 1999; Prieto and Winer, 1999; Tiong et al., 2019; Vandevelde et al., 1996). Indeed, when we injected a retrograde tracer in M1 of Ntsr1-Cre;tdTomato mice (Figure 8A), we found a clear band of retrogradely labeled deep L6 neurons in somatosensory cortex, under the layer of tdTomato-positive L6CThNs in Ntsr1-Cre;tdTomato mice, in addition to M1-projecting S1 corticocortical neurons in more superficial layers including L2/3, L5 and L6 (Figure 8B). However, whether the axons of single L6b neurons ramify locally within a cortical area as well as project to neighboring cortical areas is not clear. To determine if single S1 deep L6 neurons project to M1 as well as locally within S1, we next injected two different retrograde tracers, one in superficial S1 and one in superficial M1 (Figure 8C), and found that a substantial fraction of deep L6 neurons in S1 were double-labeled (Figure 8D,E). Thus, some L6b neurons that project locally within S1 also project to M1.

**Figure 8:**
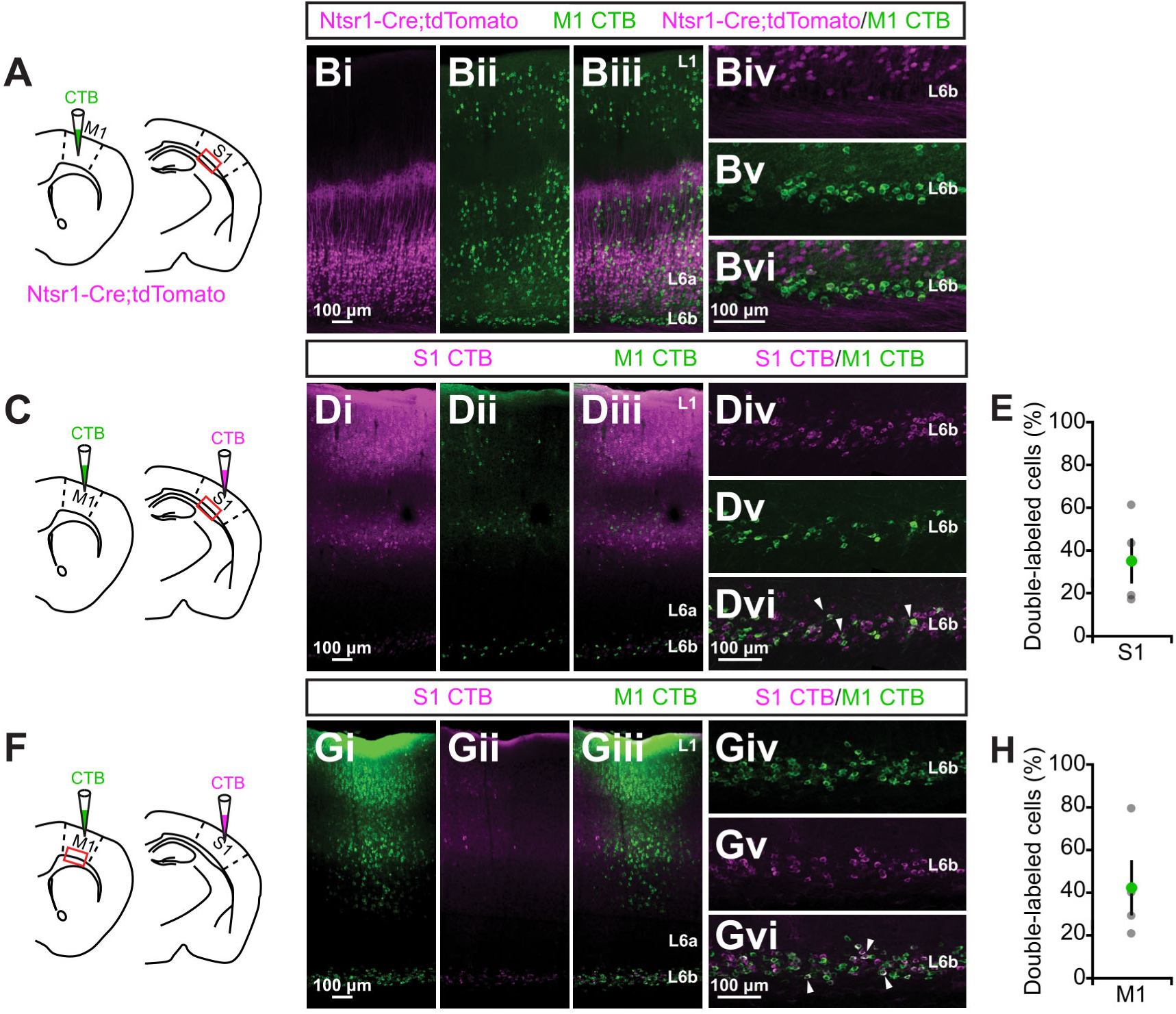
Layer 6b neurons interconnect ipsilateral cortical areas. ***A,*** Schematic showing experimental configuration. ***B,*** Images of layer 6 corticothalamic neurons (L6CThNs, magenta) in S1 of an Ntsr1-Cre;tdTomato mouse (Bi) with neurons retrogradely labeled from M1 (Bii, green, AlexaFluor 488 CTB; P47 mouse). The images are overlaid (Biii). M1-projecting deep L6 neurons (green) located below L6CThNs (magenta) in S1 are shown at higher magnification (Bv-Bvi). ***C***, Schematic showing experimental configuration. ***D,*** Images of S1 showing deep L6 neurons retrogradely labeled from layers 1 and 2 (L1/2) in S1 (magenta, AlexaFluor 488 CTB, Di) and M1 (green, AlexaFluor 647 CTB, Dii; P29 mouse). The images are overlaid (Diii). Div-Dvi show labeled neurons at higher magnification. Arrowheads indicate examples of double-labeled neurons. ***E,*** Summary data showing the percentage of L6b neurons in S1 retrogradely labeled from M1 and locally from S1 (n = 4 mice, P29-P30). ***F***, Schematic showing the experimental configuration. ***G,*** Images of M1 showing deep L6 neurons retrogradely labeled from L1/2 in M1 (Gi) and in S1 (Gii) from the same animal as in D. The two images are overlaid (Giii) and deep L6 is shown at higher magnification (Giv-Gvi). Arrowheads indicate examples of double-labeled neurons. ***H,*** Summary data showing the percentage of L6b neurons in M1 retrogradely labeled from S1 and locally from M1 (n = 4 mice, P29-P30, same mice as in E). Scale bars: B,D,G: 100 μm.

We next assessed whether deep L6 neurons in M1 project both locally in M1 as well as to S1. Again, we found double-labeled S1-projecting L6b neurons in M1 (Figure 8F-H). In these experiments, we did not identify any retrogradely labeled deep L6 neurons in the contralateral hemisphere, further confirming that neurons in deep L6 do not project to contralateral cortex. Furthermore, following bilateral injections of two retrograde neuronal tracers into the superficial layers of somatosensory cortex, we did not find any double-labeled neurons in deep L6 (n = 3 mice, P25-P30).Taken together, these data indicate that single L6b neurons in S1 and M1 ramify locally as well form long-range ipsilateral interareal connections between M1 and S1, but do not form connections with contralateral cortex.

### The electrophysiological properties of L6b neurons are stable from early adolescence through adulthood

L6b and white matter interstitial neurons are thought to represent a persistent fraction of subplate neurons (SPNs), the first neurons to be generated in the neocortex. SPNs are thought to undergo programmed cell death during the first postnatal weeks, (Allendoerfer and Shatz, 1994; Hoerder-Suabedissen and Molnár, 2013; Kanold and Luhmann, 2010; Kostovic and Rakic, 1990; Luskin and Shatz, 1985; Marx et al., 2017; Price et al., 1997; Robertson et al., 2000; Torres-Reveron and Friedlander, 2007; Woo et al., 1991; Wood et al., 1992) although some studies have found little evidence for programmed cell death (Chang et al., 2024; Ghezzi et al., 2021; Nicolelis et al., 1991; Valverde et al., 1995). As SPNs are thought to be molecularly and morphologically diverse (Hoerder-Suabedissen and Molnár, 2013; Hoerder-Suabedissen et al., 2009; Marx and Feldmeyer, 2013), with some molecular types more resistant to apoptosis than others (Hoerder-Suabedissen and Molnár, 2013), one possibility is that CTGF-positive L6b neurons represent a population that is relatively resistant to apoptosis, thus representing an increasing fraction of surviving neurons as development progresses. Alternatively, the proportion of CTGF-positive neurons in deep L6 may remain relatively constant or decrease as development progresses. To test these hypotheses, we immunostained sections from the somatosensory cortex of Ntsr1-Cre;tdTomato mice for CTGF as well as for NeuN (Hrnbp3), a marker of neuronal nuclei, to determine the proportion of the neuronal population in deep L6 that were CTGF-positive from postnatal day 2 (P2) to adulthood (P93) (Figure 9A). We found that the vast majority of NeuN-positive neurons below the major band of tdTomato-positive L6CThNs were CTGF-positive across all time points tested (Figure 9B). The slope of a line fitted across the data from P13 to P93 was not statistically different from 0, indicating that the proportion of CTGF-positive neurons relative to the total neuron population in deep L6 did not change significantly over this time period. Thus, the proportion of CTGF-positive neurons in deep L6 remains stable after the second postnatal week in mice.

**Figure 9:**
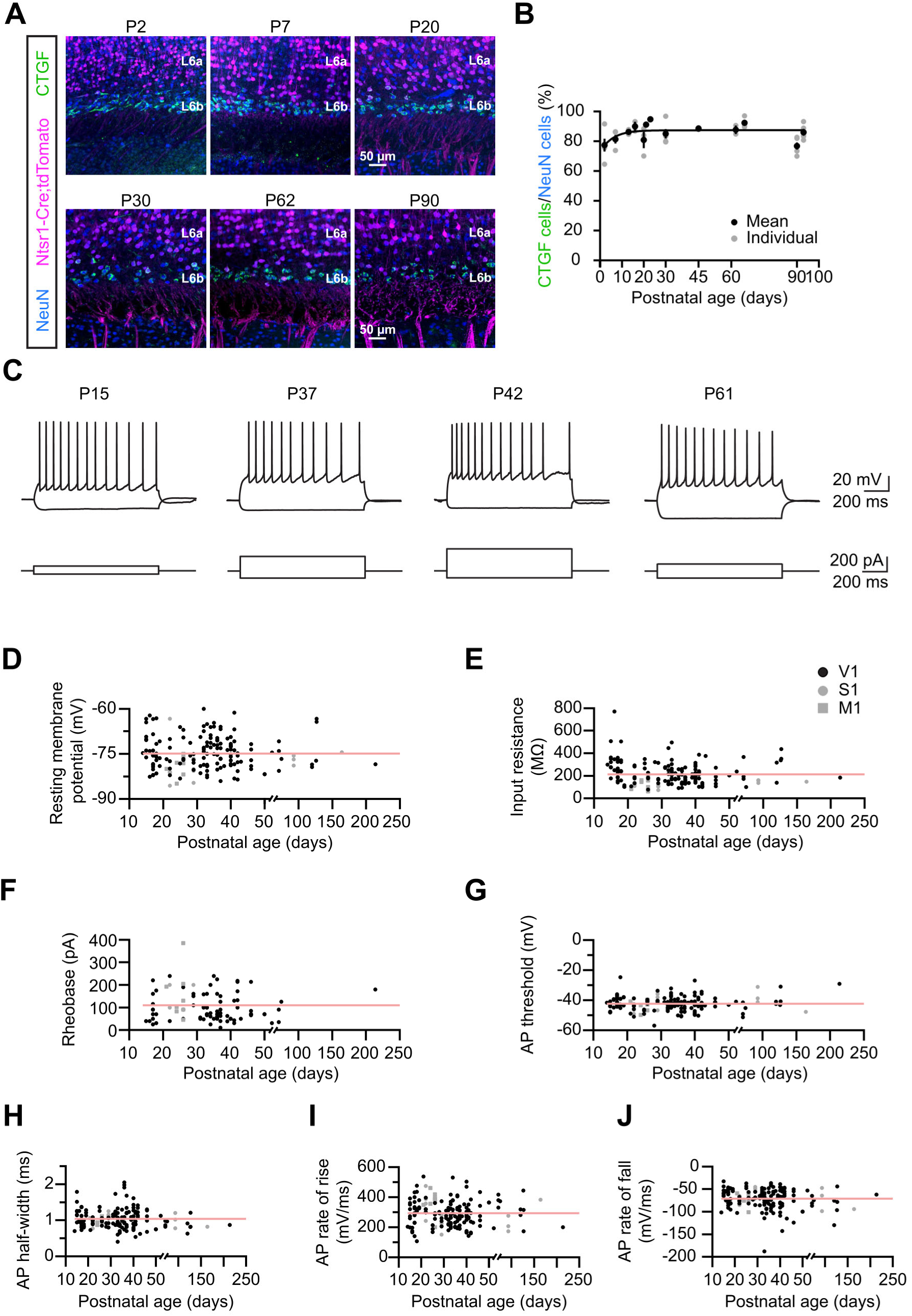
The anatomical and electrophysiological properties of L6b neurons are maintained following the first postnatal week. ***A***, Representative images of somatosensory cortex (S1) from postnatal day 2 (P2) to P90 Ntsr1-Cre;tdTomato mice showing CTGF immunostaining (green), NeuN immunostaining (blue) and tdTomato expression (magenta). ***B***, Summary data showing the proportion of CTGF-positive neurons relative to the total NeuN-positive population located beneath the tdTomato-positive L6CThNs in Ntsr1-Cre;tdTomato mice across development (P2, n = 5 mice; P7, n = 5 mice; P13, n = 2 mice; P16, n = 2 mice; P20, n = 3 mice; P21, n = 1 mouse; P23, n = 2 mice; P30, n = 5 mice; P45, n = 2 mice; P62, n = 2 mice; P66, n = 4 mice; P90, n = 5 mice; P93, n = 5 mice; Fit P13-P93: beta = –0.06, p = 0.293). ***C***, Representative voltage traces of L6b neurons in response to depolarizing and hyperpolarizing current steps from P15 to P61. The resting membrane potential (RMP, ***D***, n = 162 neurons, beta = 3.36 x 10^-5^, p = 0.998; n = 141 neurons in visual cortex, P14-P214; n = 13 neurons in S1, P22-P164; n = 8 neurons in motor cortex, P21-P26), input resistance (***E***, n = 159 neurons, beta = –0.11, p = 0.723; n = 138 neurons in visual cortex, P14-P214; n = 13 neurons in S1, P22-P164; n = 8 neurons in motor cortex, P21-P26), rheobase (***F***, n = 89 neurons, beta = 0.06, p = 0.881; n = 77 neurons in visual cortex, P15-P214; n = 4 neurons in S1, P22-P29; n = 8 neurons in motor cortex, P21-P26), action potential (AP) threshold (***G***, n = 162 neurons, beta = 0.03, p =0.053; n = 141 neurons in visual cortex, P14-P214; n = 13 neurons in S1, P22-P164; n = 8 neurons in motor cortex, P21-P26), AP half-width (***H***, n = 162 neurons, beta = –0.001, p = 0.087; n = 141 neurons in visual cortex, P14-P214; n = 13 neurons in S1, P22-P164; n = 8 neurons in motor cortex, P21-P26), AP rate of rise (***I***, n = 162 neurons, beta = –0.14, p = 0.581; n = 141 neurons in visual cortex, P14-P214; n = 13 neurons in S1, P22-P164; n = 8 neurons in motor cortex, P21-P26) and AP rate of fall (***J***, n = 162 neurons, beta = –0.10, p = 0.112; n = 141 neurons in visual cortex, P14-P214; n = 13 neurons in S1, P22-P164; n = 8 neurons in motor cortex, P21-P26) are shown. Scale bar: A: 50 μm.

Previous studies indicate that the intrinsic electrophysiological properties of SPNs vary in early development, exhibiting progressive maturation in the first two postnatal weeks (Ghezzi et al., 2021; Liao and Lee, 2012; Zhao et al., 2009). Whether the intrinsic electrophysiological properties of L6b neurons continue to evolve in juvenile and adult animals or remain stable, consistent with our anatomical findings, is less clear. To address this question, we compared the electrophysiological properties of L6b neurons recorded in acute slices of visual cortex from Ntsr1-Cre;tdTomato mice, aged P13 to P214 (Figure 9C-J). Although the majority of our recordings were from visual cortex, we also assessed the intrinsic properties of L6b neurons in somatosensory cortex and motor cortex. We found that throughout development, L6b neurons displayed mild spike frequency adaptation in response to step depolarizations and exhibited little sag response to hyperpolarizing current steps (Figure 9C). When we compared their electrophysiological properties across ages, we found no significant changes in their resting membrane potential (Figure 9D) or input resistance (Figure 9E). The rheobase values and action potential thresholds also did not significantly change after the second postnatal week (Figure 9F,G). The properties of the action potentials of deep L6 neurons were also similar from P13 to P214 (Figure 9H-J). These data indicate that the electrophysiological properties of L6b neurons are stable from 2 weeks after birth to adulthood (3-7 months).

### Layer 6b neurons in primary visual cortex are not integrated into thalamocortical circuits involving primary sensory thalamic nuclei

Subplate neurons are thought to play an important role in establishing cortical circuits during early development (Hoerder-Suabedissen and Molnár, 2015; Kanold and Luhmann, 2010; Molnár et al., 2020; Ohtaka-Maruyama, 2020). Studies in sensory cortex indicate that thalamocortical axons synapse onto SPNs in the first postnatal weeks (Friauf et al., 1990; Hanganu et al., 2002; Molnár et al., 2003; Zhao et al., 2009 although see Ghezzi et al., 2021). In contrast, relatively weak thalamocortical input onto deep L6 neurons was recently described in somatosensory cortex of adult mice (Frandolig et al., 2019; Zolnik et al., 2020), but how L6b neurons are integrated into geniculocortical circuits in adult mice remains unclear. To determine whether L6b neurons in the visual cortex of adult mice receive geniculocortical input, we expressed channelrhopdopsin-2 (ChR2) in geniculocortical axons in Ntsr1-Cre;tdTomato mice and recorded light-evoked postsynaptic potentials (PSPs) in pairs of neurons composed of one L6b neuron and one tdTomato-positive L6CThN (Figure 10A). We observed significantly weaker responses to geniculocortical input in L6b neurons than in L6CThNs (P37-45), with some L6b neurons receiving no detectable geniculocortical input (Figure 10B,C). Thus, although geniculocortical axons pass through deep L6 towards their more pial targets, L6b neurons receive little geniculocortical input in adult mice.

**Figure 10:**
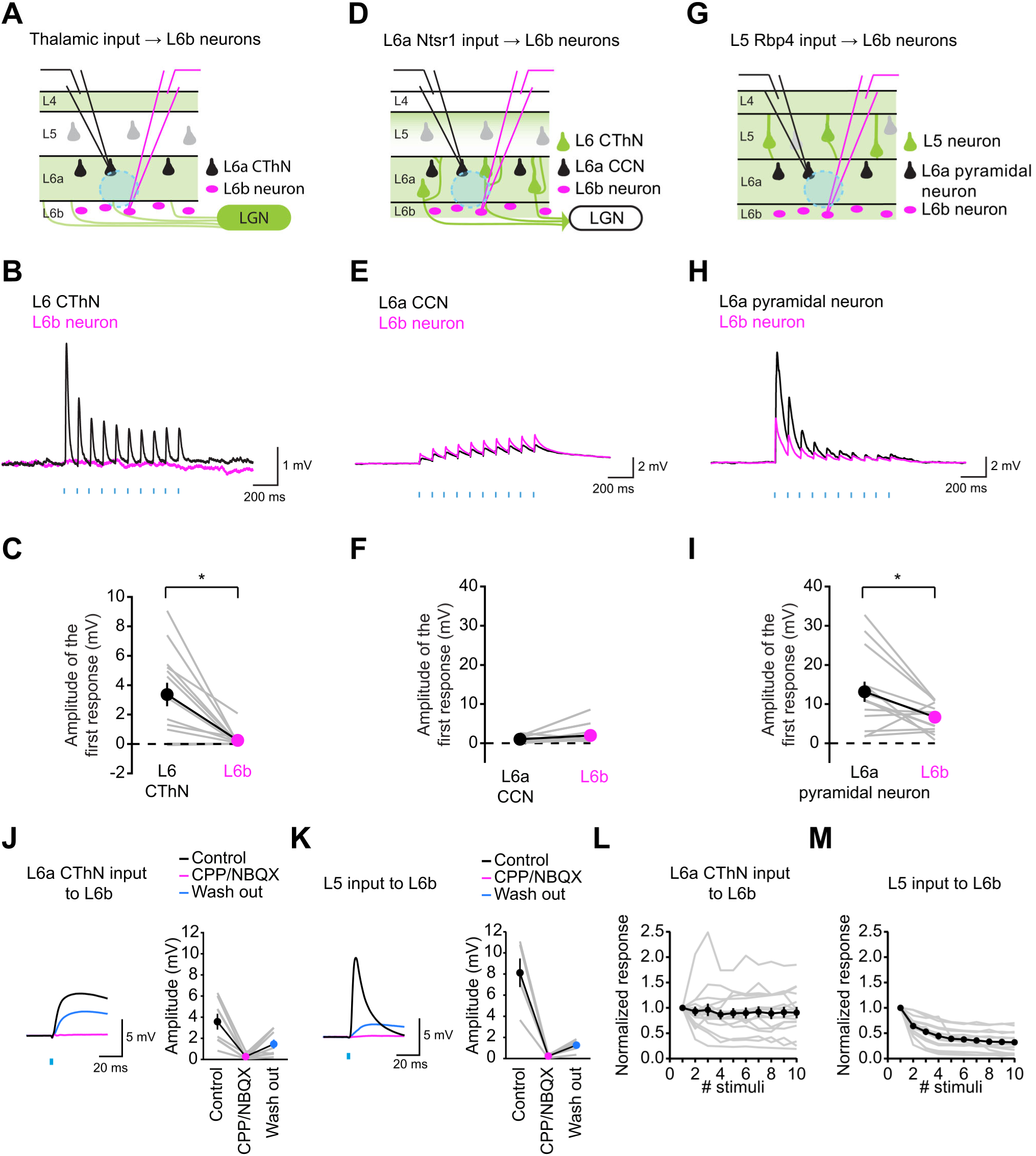
L6b neurons in visual cortex in adult mice receive little geniculocortical input. ***A***, Experimental configuration for recording responses to optogenetic stimulation of geniculocortical fibers in pairs of neurons composed of one L6a corticothalamic (CThN) neuron and one L6b neuron from the visual cortex of Ntsr1-Cre;tdTomato mice. ***B***, Example responses (postnatal day 45 (P45) mouse). ***C***, Summary data of the amplitudes of the first response for pairs composed of a L6b neuron and a L6CThN recorded in visual cortex (n = 13 pairs from 3 mice, P37-P45, Wilcoxon signed rank test, p = 0.014). ***D***, Experimental configuration for recording responses to optogenetic stimulation of L6CThN neurons in pairs composed of one L6b neuron and one L6a corticocortical neuron from the visual cortex of Ntsr1-Cre;ChR2-YFP mice. ***E***, Example responses (P32 mouse). ***F***, Summary data of the amplitudes of the first response (n = 11 pairs from 6 mice, P26-P40, Wilcoxon signed rank test, p = 0.652). ***G***, Experimental configuration for recording responses to optogenetic stimulation of L5 neurons in pairs composed of one L6b neuron and one L6a neuron in the visual cortex of Rbp4-Cre;ChR2-YFP mice. ***H***, Example responses (P34 mouse). ***I***, Summary data of the amplitudes of the first response for pairs of L6b and L6a neurons in visual cortex of Rbp4-Cre;ChR2-YFP mice (n = 14 pairs from 6 mice, P31-P42, Wilcoxon signed rank test, p = 0.044). ***J***, Representative voltage traces recorded from a L6b neuron in response to photostimulation of L6CThN input before (black), during (red), and after (blue) bath application of glutamate receptor antagonists (left, P39 mouse) and summary data (right; n = 8 neurons from 6 mice, P33-P41). ***K***, Representative voltage traces recorded from a L6b neuron in response to photostimulation of L5 Rbp4 input before (black), during (red), and after (blue) bath application of glutamate receptor antagonists (left, P33), and summary data (right; n = 5 neurons from 3 mice, P38-P42). ***L***, Summary data showing responses in L6b neurons to 10 Hz photostimulation of L6CThN input (Ntsr1-Cre;ChR2-YFP) normalized to the amplitude of the first response (n = 16 neurons from 8 mice, P32-P40). Individual responses are shown in grey and the mean in black. ***M***, Summary data showing normalized amplitudes of responses in L6b neurons to 10 Hz photostimulation of L5 Rbp4-Cre;ChR2-YFP input (n = 14 neurons from 6 mice, P31-P42).

We next asked whether L6b neurons in visual cortex receive input from L6CThNs. Not only do the axons of L6CThNs traverse L6b on their way to the thalamus, but the dendrites of L6b neurons also ramify widely throughout L6 (Figure 5A,B). We recorded light-evoked responses from neuron pairs composed of one L6b neuron and one ChR2-negative L6 corticocortical

neuron (CCN) in slices of visual cortex from Ntsr1-Cre;ChR2-YFP mice. (Figure 10D). We recorded only weak PSPs in CCNs, consistent with studies showing that L6CThNs rarely synapse onto L6 CCNs (Figure 10E,F) (Crandall et al., 2017; West et al., 2006). Responses in L6b neurons were similarly weak. Using the same approach with a L5 Cre line, Rbp4-Cre (Gerfen et al., 2013), we found that both L6b neurons and L6a neurons responded robustly to activation of L5 neurons, although the amplitudes of the responses were, on average, significantly smaller in L6b neurons relative to neurons in upper L6 (Figure 10G-I). The L6b responses to both L6CThN and L5 input were primarily mediated by glutamate receptors as bath application of antagonists to AMPA and NMDA receptors abolished the responses (Figure 10J,K). However, the short-term synaptic dynamics of the two connection types were significantly different (Figure 10L,M; Two-way ANOVA, Input type: p = 4.67e-26; Stimulus number: 5.82e-05; Input type x Stimulus number: p = 0.018). Although differences in the synaptic properties tested using ChR2 must be interpreted with caution (Jackman et al., 2014), the responses to L5 input consistently showed strong depression in contrast to the L6CThN responses, which showed more variable short-term synaptic plasticity, ranging from substantial facilitation to strong depression. Taken together, these data indicate that while L6b neurons in visual cortex are embedded into cortical circuits, they are not strongly integrated into geniculocortical or layer 6 corticothalamic circuits in the visual cortex of adult mice.

## Discussion

### Consensus definition for L6b neurons

By quantitatively comparing different methods used to denote neurons in the deepest layer of the neocortex, termed L6b neurons, we identified several methods that defined similar neuron populations, generating a consensus definition for L6b neurons. Injecting retrograde neuronal tracers in the superficial layers of the cortex reliably labeled a thin band of neurons in the bottom 10% of the somatosensory cortex, consistent with work in the rat (Clancy and Cauller, 1999).

This cell population was largely immunopositive for CTGF and Cplx3, although immunostaining for CTGF may capture a slightly larger and Cplx3 a slightly smaller overall cell population. We also found that this population of deep L6 neurons retrogradely labeled from superficial cortical layers was primarily located between the white matter border and Cre-expressing L6CThNs in the Ntsr1-Cre mouse line, although some L6b neurons express Cre recombinase in these mice. We found that Cre expression in the transgenic mouse line, Nxph4-CreER also identified this neuron population. Of the markers tested, *Nxph4* was most restricted to L6b neurons in neocortex although neuron classes outside the neocortex also express *Nphx4*, some of which may project into the neocortex, an important consideration when genetically accessing this cortical cell population. We found that almost none of the neurons defined using this consensus definition were labeled following injections of retrograde neuronal tracers into somatosensory or visual thalamus, indicating that this cell population in S1 and V1 does not include corticothalamic neurons.

The neurons in this cell population were excitatory neurons with spiny, multipolar dendritic arbors extending throughout L6. Although prior studies have highlighted a diversity of cell types in deep layer 6 using a variety of techniques (Andjelic et al., 2009; Arimatsu et al., 2003; Marx and Feldmeyer, 2013; Marx et al., 2017; Scala et al., 2021; Tasic et al., 2018; Yao et al., 2021; Zolnik et al., 2024; Zolnik et al., 2020), we found that L6b neurons identified using this consensus definition had similar dendritic morphologies. None of the neurons we filled exhibited a clear apical dendrite. Thus, we propose that the neuron population in deep L6 that is retrogradely labeled from L1/2, expresses CTGF, Cplx3 and Nxph4 and does not project to the thalamus be designated L6b neurons. These neurons represent the dominant excitatory neuron type in L6b. However, differences in molecular marker expression may exist across cortical areas (Figures 2, 3F). For example, earlier in development, CTGF and Nxph4 appear expressed in a smaller fraction of neurons in primary auditory cortex than in primary somatosensory and visual cortex while Cplx3 is expressed more similarly across these cortical areas (Chang et al., 2024). Future work will further define how the molecular identity of L6b neurons varies across cortical areas and will test whether molecular subtypes relate to functional subtypes of excitatory L6b neurons.

### Cre expression in Drd1a-Cre mice captures heterogeneous subpopulations of neurons

In contrast to the limited vertical distribution of L6b neurons identified using our consensus definition, tdTomato-positive neurons in Drd1a-Cre;tdTomato mice were observed throughout the vertical extent of L6. The dendritic morphology of individual tdTomato-positive neurons in the somatosensory cortex of Drd1a-Cre;tdTomato mice was consistent with these findings: some neurons in deep layer 6 were spiny, multipolar cells similar to L6b neurons. However, the majority of tdTomato-positive cells that we filled in acute slices of somatosensory cortex from Drd1a-Cre;tdTomato mice were pyramidal neurons with prominent apical dendrites extending into at least L5a, similar in morphology to other L6 cell types including L6CThNs projecting to POm (Frandolig et al., 2019; Zhang and Deschênes, 1997). The morphological diversity of Cre-expressing neurons in the cortex of Drd1a-Cre mice is also consistent with our scRNA-seq analyses showing *Drd1* expression in neurons annotated as L6b neurons, L6CThNs, L6 IT neurons and L5/6 non-projecting pyramidal neurons. These findings are also consistent with studies showing that POm-projecting Drd1a-Cre neurons in S1 send axons to L5a, as expected for POm-projecting L6CThNs, while the axons of Drd1a-Cre neurons transfected from L1 do not ramify in L5a but rather L1 (Chevée et al., 2018; Frandolig et al., 2019; Zhang and Deschênes, 1997; Zolnik et al., 2024).

In addition, neither the L6b nor the POm-projecting L6CThN populations were fully captured by Cre expression in the somatosensory cortex of Drd1a-Cre;tdTomato mice. Less than 20% of retrogradely labeled L6b neurons were tdTomato-positive, and only approximately 30% of retrogradely labeled L6CThNs projecting to POm were tdTomato-positive. Furthermore, both of the cell populations represent only a fraction of the Cre-expressing neurons in Drd1a-Cre;tdTomato mice: approximately 6% of Cre expressing neurons represent the subset of L6b neurons expressing Cre (< 20% of L6b neurons) and approximately a quarter of Cre expressing neurons represent the Cre-expressing L6CThNs (∼30% of L6CThNs) projecting to higher order thalamic nuclei. Thus, as suggested in previous studies (Ansorge et al., 2020; Hoerder-Suabedissen et al., 2018; Zolnik et al., 2024; Zolnik et al., 2020), Cre is expressed in a heterogeneous population of neurons in Drd1a-Cre mice, many of which are neither L6b neurons nor L6CThNs. Together, these results indicate that the influence of deep layer 6 neurons on thalamic function in studies using Drd1a-Cre mice is likely due to Cre-expressing L6CThNs projecting to higher order thalamic nuclei rather than to L6b neurons identified using our consensus definition (Ansorge et al., 2020; Hoerder-Suabedissen et al., 2018; Zolnik et al., 2024). Why subsets of neurons in deep L6 express Cre in Drd1a-Cre mice is not clear. Whether these Cre-expressing neurons capture specific subtypes of L6b neurons and L6CThNs or whether Cre-expressing neurons are in different cell states, for example, awaits further study.

### L6b neurons in adult mice are not integrated into thalamocortical circuits of primary sensory thalamic nuclei

Using the consensus definition, we showed that L6b neurons in visual and somatosensory cortex do not project to the thalamus. In addition, we found that L6b neurons in visual cortex receive little geniculocortical input, in contrast to the significant thalamocortical input to subplate neurons in visual cortex in early development (Friauf et al., 1990; Friauf and Shatz, 1991). Our findings in visual cortex are consistent with work in somatosensory cortex showing little anatomical or functional thalamocortical input to neurons in deep L6 in adult mice (Frandolig et al., 2019; Meyer et al., 2010; Wimmer et al., 2010; Zolnik et al., 2020), in contrast to the significant thalamocortical input onto subplate neurons in the period before thalamocortical axons have reached L4 in sensory cortical areas (Hanganu et al., 2002; Higashi et al., 2002; Kanold, 2009; Molnár et al., 2003) although see (Ghezzi et al., 2021). In addition, as has been previously reported for S1 (Zolnik et al., 2020), we found that L6b neurons in visual cortex received little input from L6CThNs. Together, these results indicate that L6b neurons are not strongly integrated into thalamocortical circuits related to primary sensory thalamic nuclei and do not play a prominent direct role in these thalamocortical interactions: they neither project to the thalamus nor receive strong thalamocortical or corticothalamic input. Together, these results also suggest that, although subplate neurons in early development and L6b neurons in adult mice exhibit responses to external sensory signals that can be sculpted by sensory experience (Meng et al., 2021; Mukherjee and Kanold, 2022; Mukherjee et al., 2023; Wess et al., 2017; Yoneda et al., 2023), these responses must be mediated by different neuronal circuits, circuits that may remodel during early development.

### L6b neurons interconnect ipsilateral cortical areas but do not project callosally

Another inconsistency across studies has been whether L6b neurons project to the contralateral cortex (Clancy and Cauller, 1999; Hoerder-Suabedissen et al., 2018; Ledderose et al., 2023; Tiong et al., 2019). We found that L6b neurons defined using the consensus definition did not project to contralateral cortex. Our results are consistent with other studies using retrograde neuronal tracers that failed to identify L6b neurons projecting to contralateral cortex (Clancy and Cauller, 1999; Ledderose et al., 2023; Tiong et al., 2019). The callosal projections identified using Drd1a-Cre mice (Hoerder-Suabedissen et al., 2018) are consistent with our scRNA-Seq analyses indicating that *Drd1* is expressed in L6 intratelencephalic (IT) neurons. Rather than projecting to cortical areas via the corpus callosum, L6b neurons interconnected ipsilateral cortical areas (Arimatsu et al., 2003; Clancy and Cauller, 1999). Indeed, a large fraction of L6b neurons were double-labeled following retrograde tracer injections into somatosensory and motor cortex, indicating that individual L6b neurons project both locally and to distant ipsilateral cortical areas and are positioned to influence cortical activity within a cortical area as well as between cortical regions.

### Layer 6b neurons exhibit consistent properties from juvenile to adult mice

Neurons in L6b are thought to originate from subplate neurons (SPNs), the first cortical neurons generated during early development (Allendoerfer and Shatz, 1994; Hoerder-Suabedissen and Molnár, 2015). Studies indicate that SPNs represent a morphologically and electrophysiologically heterogeneous neuronal population that undergoes programmed cell death with a small fraction persisting to become L6b neurons or interstitial white matter neurons (Chun and Shatz, 1989; Hoerder-Suabedissen and Molnár, 2013; Kostovic and Rakic, 1990; Luskin and Shatz, 1985; Marx and Feldmeyer, 2013; Marx et al., 2017; Price et al., 1997; Robertson et al., 2000; Torres-Reveron and Friedlander, 2007; Woo et al., 1991; Wood et al., 1992), although some studies have found little evidence for programmed cell death (Chang et al., 2024; Ghezzi et al., 2021; Nicolelis et al., 1991; Valverde et al., 1995). We found that the fraction of L6b neurons in deep L6 identified using our consensus definition remained consistent from approximately P14 to adulthood (∼3 months of age). Further, we found that the intrinsic electrophysiological properties of L6b neurons were established by P14 and remained similar through adulthood (> 3 months old). Thus, although the cortex continues to mature after P14, our results indicate that the basic morphological and electrophysiological properties of L6b neurons are established by this time point.

### Proposed nomenclature for layer 6 of the rodent neocortex

Based on these data, we propose that L6 in somatosensory and visual cortex of adult rodents be divided into two major sublayers. We propose that the thin layer, primarily composed of multipolar excitatory neurons that are retrogradely labeled from superficial cortical layers and are largely CTGF-positive, Cplx3-positive and Nxph4-positive, be designated L6b. We propose that the overlying layer containing L6CThNs as well as other pyramidal cell types be designated L6a. Data from our lab and others indicate that, in rodents, L6a can be further subdivided based on the axonal project patterns of L6CThNs. The cell bodies of L6CThNs that project only to primary sensory thalamic nuclei like LGN and VPM are biased to upper L6a (Bourassa and Deschênes, 1995; Bourassa et al., 1995; Chevée et al., 2018; Frandolig et al., 2019; Whilden et al., 2021). This sublayer of L6a also includes interlaminar parvalbumin-positive interneurons (Frandolig et al., 2019). The cell bodies of L6CThNs that project to higher order thalamic nuclei like LP and POm are biased towards lower L6a (Bourassa and Deschênes, 1995; Bourassa et al., 1995; Chevée et al., 2018; Frandolig et al., 2019; Whilden et al., 2021) and may include L6CThNs that project to both first order and higher order thalamic nuclei (i.e. VPM/POm-projecting and LGN/LP-projecting L6CThNs) as well as L6CThNs that project to POm or LP only (Bourassa and Deschênes, 1995; Bourassa et al., 1995; Hoerder-Suabedissen et al., 2018).

This latter population appears to be the main L6CThN population expressing Cre in Drd1a-Cre mice as few axons are seen in LGN or VPM in Drd1a-Cre;tdTomato mice (Hoerder-Suabedissen et al., 2018). Furthermore, their dendritic morphology is also consistent with this interpretation (Hoerder-Suabedissen et al., 2018; Zhang and Deschênes, 1997). Although the features of L6b neurons that we assessed were similar in M1, how generalizable this overall organization is across cortical areas remains to be tested. For example, how L6CThN projections and L6b neurons are organized in the auditory system remains to be fully elucidated (Bajo and King, 2023). Using clear, shared working definitions of the cell types found in L6 will precisely enable such comparisons across studies of the synaptic organization of these cell types, their contributions to cortical activity and their influence on perception and behavior.

## Materials and Methods

### Mice

All experimental procedures were approved by the Johns Hopkins Animal Care and Use Committee and conducted in accordance with the guidelines of the National Institutes of Health and the Society for Neuroscience. The following mouse lines were used: Neurotensin receptor-1 Cre (Ntsr1-Cre, GENSAT 220; RRID: MMRRC_030648-UCD) (Gong et al., 2007),

Neurexophilin 4 2A-CreERT2-D (Nxph4-CreER, Jackson Laboratories 022861, RRID:IMSR_JAX:022861) (Harris et al., 2014), Retinol binding protein-4-Cre (Rbp4-Cre, GENSAT KL100, RRID:MMRRC_031125-UCD) (Gerfen et al., 2013), Dopamine receptor 1a-Cre (Drd1a-Cre, GENSAT FK164, RRID:MMRRC_030781-UCD, a gift of Hongkui Zeng) (Gerfen et al., 2013), Gad2-T2a-NLS-mCherry (Gad2-mCherry, Jackson Laboratories 023140, RRID:IMSR_JAX:023140) (Peron et al., 2015), loxP-STOP-loxP-tdTomato Cre reporter (Ai9 and Ai14, Allen Institute for Brain Science, RRID:IMSR_JAX:007905, RRID:IMSR_JAX:007908) (Madisen et al., 2010), and a mouse line carrying a Cre-dependent channelrhodopsin-2-YFP construct (Ai32, Allen Institute for Brain Science, RRID:IMSR_JAX:024109) (Madisen et al., 2012). All lines were maintained on a mixed background composed primarily of C57BL/6J and CD-1, and mice of both sexes were used for experiments. All animals were maintained on a 12 h light/dark cycle with *ad libitum* access to food and water.

### Stereotaxic injections of retrograde neuronal tracers and viral constructs

For stereotaxic injections of retrograde neuronal tracers and viruses, mice of either sex were first anesthetized with ketamine (50 mg/kg), dexmedetomidine (25 µg/kg) and the inhalation anesthetic, isoflurane. Animals were placed in a custom-built stereotaxic frame and anesthesia was maintained with isoflurane (1.5-2 %). A small craniotomy was performed, a glass pipette (12.5-35 µm tip diameter) loaded with tracer or virus was lowered into the brain, and the material was pressure-injected. Following the injection, the pipette was retracted, the incision was sutured and the analgesic, buprenorphine (0.05 mg/kg), was administered to all animals postoperatively.

To retrogradely label deep L6 neurons, 45-90 nl of fluorophore-conjugated (AlexaFluor 488, 555 or 647) cholera toxin subunit B (CTB; C34775, C34776 or C34778, ThermoFisher Scientific) or 45-90 nl of red Retrobeads or Retrobeads IX (Lumafluor) were injected into either primary somatosensory cortex (S1, AP: –1 mm, ML: 2.85 mm, 0.15 mm from the surface of the cortex), primary visual cortex (V1, AP: 3.45 mm, ML: 2.7 mm, 0.15 mm from the surface of the cortex) or primary motor cortex (M1, AP: 0.9 mm, ML: 1.45 mm, 0.15 mm from the surface of the cortex). To retrogradely label corticothalamic and corticocortical neurons, 45-90 nl of CTB were injected into the posterior nucleus of the thalamus (PO, AP: –2 mm, ML: 1.2 mm DV: 3 mm), the lateral posterior nucleus of the thalamus (LP, AP: –2 mm, ML: 1.7 mm, DV: 2.5 mm), or contralateral S1 (cS1, AP: – 1 mm, ML: –2.85 mm, DV: 0.5 mm and 1 mm). Mice were sacrificed 4-7 days following the injections.

To express channelrhodopsin-2 (ChR2) in geniculocortical neurons, 30-90 nl of AAV-DJ-CaMKIIα-ChR2-eYFP (A gift of Karl Deisseroth, GVVC-AAV-36, 1.90E+13 vg/mL, Stanford Gene Vector and Virus Core) was injected into the LGN (AP: 2.25 mm, ML: 2.4 mm, DV: 3.1 mm) of postnatal day 20-31 (P20-P31) Ntsr1-Cre;tdTomato mice or C57BL/6J mice with L6b neurons retrogradely labeled from L1/2. Mice were sacrificed 14-18 days after virus injection. To test the synaptic input from layer 5 pyramids or layer 6 corticothalamic neurons onto deep L6 neurons in visual cortex, Rbp4-Cre;ChR2-YFP or Ntsr1-Cre;ChR2-YFP mice were used, and 45-90 nl AlexaFluor 555-conjugated CTB or red Retrobeads or Retrobeads IX (Lumafluor) were injected into L1/2 of the visual cortex as described above.

### Tamoxifen induction of Cre expression in Nxph4-CreER mice

To induce Cre expression in Nxph4-CreER mice, tamoxifen (Sigma, T5648) was dissolved in sunflower seed oil (Sigma, S5007) at a concentration of 10 mg/ml. Nxph4-CreER mice (P18-P108) were injected intraperitoneally with tamoxifen (100 mg/kg) for 5 consecutive days. To induce Cre expression in pups starting at P4, tamoxifen was administered to dams using the same protocol as above. Mice injected with tamoxifen for the expression of Cre recombinase were sacrificed two or more days after the last tamoxifen injection.

### Immunohistochemistry

Table 1 includes a summary of the antibodies used. To compare the distribution of CTGF-positive neurons relative to retrogradely labeled deep L6 neurons and tdTomato-positive neurons in Ntsr1-Cre;tdTomato and Nxph4-CreER;tdTomato, mice were perfused transcardially with 10 mL of 0.01 M phosphate buffered saline (PBS) followed by 10 ml of 4% paraformaldehyde (PFA, Electron Microscopy Sciences, 15710) in PBS. The brain was then removed and postfixed for at least 3 hours in 4% PFA in 0.01 M PBS. The brain was hemisected, hemispheres mounted on a 30° ramp, and 50 µm or 100 µm-thick parasagittal sections were cut on a vibratome (VT-1000S, Leica) before being subjected to immunohistochemistry. Sections were first incubated for 1 h in blocking solution (0.01 M PBS with 0.3% Triton X-100, Sigma, T9284; 5% normal donkey serum, NDS, VWR, 80054-446 or Calbiochem, 566460-5ML) at room temperature, then incubated with antibodies to CTGF (goat anti-CTGF, 1:300, Santa Cruz Biotechnology, SC-14939), to DsRed (Rabbit anti-DsRed, 1:1000, 632496, Takara Bio) and/or to NeuN (mouse anti-NeuN, 1:500, MAB377, Millipore) in PBS with 0.3% Triton X-100 and 5% NDS for at least 24 hours at room temperature. In some experiments, an alternative antibody to CTGF was used (mouse anti-CTGF, 1:300, Santa Cruz Biotechnology, SC-365970). This second anti-CTGF antibody required an antibody retrieval process with citric acid buffer (10 mM citric acid, pH 6.0, Sigma, 251275; 0.05% Tween 20, Sigma, P1379) performed for 30 min at 95°C prior to the initial blocking step. Sections were then rinsed, typically 3x for 10 min, in 0.01 M PBS, and then incubated with the corresponding secondary antibodies (AlexaFluor 488-conjugated Donkey anti-Goat IgG (H+L) antibody, 1:300, A11055, ThermoFisher Scientific; AlexaFluor 594-conjugated Donkey anti-Rabbit antibody, 1:1000, R37119, ThermoFisher Scientific; AlexaFluor 647-conjugated Donkey anti-Mouse antibody, 1:1,000, A31571, ThermoFisher Scientific) for at least 12 hours at room temperature. Sections were then mounted in Aqua-Poly/Mount (Polysciences Inc., 18606-20).

**Table 1:** List of antibodies used in experiments.

| **Antibody** | **Source** | **Catalog number and identifier** |
| --- | --- | --- |
| Chicken anti-GFP antibody | Aves | Cat. No.: GFP-1020; RRID: AB_10000240 |
| Rabbit anti-DsRed antibody | Takara Bio | Cat. No.: 632496; RRID: AB_10013483 |
| Goat anti-Connective Tissue Growth Factor (CTGF) antibody | Santa Cruz Biotechnology | Cat. No.: SC-14939; RRID: AB_638805 |
| Mouse anti-Connective Tissue Growth Factor (CTGF) antibody | Santa Cruz Biotechnology | Cat. No.: SC-365970; RRID: AB_10917259 |
| Rabbit anti-Complexin 3 antibody | Synaptic Systems | Cat. No.: 122 302; RRID: AB_2281240 |
| Mouse anti-NeuN antibody | Millipore | Cat. No.: MAB377; RRID: AB_2298772 |
| Goat anti-Choline Acetyltransferase (ChAT) antibody | Millipore | Cat. No.: AB144; RRID: AB_90650 |
| Goat anti-Orexin-A antibody | Santa Cruz Biotechnology | Cat. No.: SC-8070; RRID: AB_653610 |
| Rabbit anti-Tryptophan hydroxylase 2 antibody | Abcam | Cat. No.: ab111828; RRID: AB_10862137 |
| Rabbit anti-Parvalbumin antibody | Swant | Cat. No.: PV27; RRID: AB_2631173 |
| Rabbit anti-Somatostatin-14 antibody | BMA biomedicals | Cat. No.: T-4103; RRID: AB_518614 |
| Chicken anti-Calbindin D-28K antibody | Novus Biologicals | Cat. No.: NBP2-50028 |
| AlexaFluor 647-conjugated Donkey anti-Rabbit IgG (H+L) antibody | Jackson ImmunoResearch Labs | Cat. No.: 711-606-152; RRID: AB_2340625 |
| AlexaFluor 488-conjugated Donkey anti-Chicken IgY (H+L) antibody | Jackson ImmunoResearch Labs | Cat. No.: 703-545-155; RRID: AB_2340375 |
| AlexaFluor 594-conjugated Donkey anti-Rabbit IgG (H+L) antibody | ThermoFisher Scientific | Cat No.: R37119; RRID: AB_141637 |
| AlexaFluor 488-conjugated Donkey anti-Rabbit IgG (H+L) antibody | ThermoFisher Scientific | Cat No.: A-21206; RRID: AB_253579 |
| AlexaFluor 488-conjugated Donkey anti-Goat IgG (H+L) antibody | ThermoFisher Scientific | Cat No.: A-11055; RRID: AB_2534102 |
| AlexaFluor 647-conjugated Donkey anti-Mouse IgG (H+L) antibody | ThermoFisher Scientific | Cat No.: A-31571; RRID: AB_162542 |
| AlexaFluor 594-conjugated Donkey anti-Mouse IgG (H+L) antibody | ThermoFisher Scientific | Cat No.: A-21203; RRID: |
| AlexaFluor 488-conjugated Donkey anti-Rabbit IgG (H+L) antibody | ThermoFisher Scientific | Cat No.: R371188; RRID: AB_2556546 |
| AlexaFluor 647-conjugated Donkey anti-Guinea pig IgG (H+L) antibody | Jackson ImmunoResearch Labs | Cat. No.: 706-605-148; RRID: AB_2340476 |

To immunostain for Cplx3, mice were perfused, and brains fixed as described above. Sections, 100 μm thick, were cut on a VT1000S coronally, incubated for 1 h in 0.01 M PBS with 0.3% Triton X-100 and 5% NDS at room temperature, and then incubated with antibodies to Cplx3 (rabbit anti-Complexin 3, 1:500, 122 302, Synaptic Systems). After rinsing, typically 3x 10 min in 0.01 PBS, sections were incubated in secondary antibodies (AlexaFluor 488-conjugated Donkey anti-Rabbit IgG (H+L) antibody, 1:1000, R37118, ThermoFisher Scientific; or AlexaFluor 594-conjugated Donkey anti-Rabbit antibody, 1:1000, R37119, ThermoFisher Scientific) for 1-2 h at room temperature. Sections were then mounted in Aqua-Poly/Mount.

To characterize Nxph4-CreER mice, mice were perfused, and brains fixed as described above. Each brain was cut parasagittally or coronally on a vibratome (VT-1000S, Leica) as described above. Sections were incubated for 1 h in 0.01 M PBS with 0.3% Triton X-100 and 3% NDS at room temperature. Slices were then incubated with primary antibodies overnight at 4°C (Goat anti-Choline acetyltransferase (ChAT, 1:200 Millipore, AB144), Goat anti-Orexin-A (1:400, Santa Cruz, SC-8070), Chicken anti-Calbindin D28K (Novus Biologicals, NBP2-50028), or Rabbit anti-Tryptophan hydroxylase 2 (TPH2, 1:400, Abcam, ab111828)). After rinsing as described above, sections were incubated with corresponding secondary antibodies (AlexaFluor 488-conjugated Donkey anti-Goat IgG (H+L) antibody, 1:300, A11055, ThermoFisher Scientific; AlexaFluor 488-conjugated Donkey anti-Rabbit antibody, 1:300, R371188, ThermoFisher Scientific; AlexaFluor 488-conjugated Donkey anti-Chicken antibody, 1:300, 703-545-155, Jackson ImmunoResearch Labs). Images were acquired with 5x (0.15 N.A.), 10x (0.3 N.A.) or 20x (0.8 N.A.) objectives on a confocal microscope (LSM 800, Zeiss) or on an epifluorescence microscope (Keyence BZ-X 710) using a 4x (0.20 N.A.) objective lens.

For the analysis of CTGF co-expression with parvalbumin or somatostatin, mice were perfused, and brains fixed as described above. After fixation, brains were cryoprotected in 30% (w/v) sucrose in PBS for 24 h. The brains were then embedded in an OCT compound and sectioned into 50 μm-thick slices using a cryostat (Leica, CM1860). The following antibodies were used for immunohistochemistry: mouse anti-CTGF antibody (1:100, Santa Cruz, SC-365970) and rabbit anti-Parvalbumin antibody (1:1,000, Swant, PV27), or rabbit anti-Somatostatin-14 antibody (1:500, BMA biomedicals, T-4103), followed by secondary antibodies: AlexaFluor 594-conjugated Donkey anti-Mouse antibody (1:300, ThermoFisher Scientific, A-21203) and AlexaFluor 488-conjugated Donkey anti-Rabbit antibody (1:300, ThermoFisher Scientific, A-21206). Images were acquired with 10x (0.45 N.A.) or 40x (1.15 N.A.) objectives on a confocal microscope (Andor Technology, Dragonfly 502w).

### Brain slice preparation and cell type identification for patch clamp recordings

To fill individual neurons with biocytin to assess their morphology and to profile the electrophysiological and synaptic properties of recorded neurons, mice (P14-P214) were anesthetized with isoflurane, and their brains rapidly removed in an ice-cold sucrose solution containing the following (in mM): 76 NaCl, 25 NaHCO3, 25 glucose, 75 sucrose, 2.5 KCl, 1.25 NaH_2_PO_4_, 0.5 CaCl_2_, 7 MgSO_4_, pH 7.3, 315 mOsm. The brain was hemisected along the midline, and one hemisphere was mounted on a 30° ramp. Acute parasagittal slices of visual cortex or somatosensory cortex, 300 μm thick, were then sectioned in the same ice-cold sucrose cutting solution using a vibratome (VT-1200s, Leica). For sections from motor cortex, a coronal cut was used to remove the anterior region of the brain, brains were then mounted on the cut face tilted anteriorly at a 15° angle, and acute brain slices (300 μm) were prepared.

Slices were incubated in warm (32-35°C) sucrose solution for approximately 30 min and then transferred to warm (32-35°C) artificial cerebrospinal fluid (aCSF) composed of the following (in mM): 125 NaCl, 26 NaHCO_3_, 2.5 KCl, 1.25 NaH_2_PO_4_, 1 MgSO_4_, 20 D-(+)-glucose, 2 CaCl_2_, 0.4 ascorbic acid, 2 pyruvic acid, 4 L-lactic acid, pH 7.3, 315 mOsm. Slices were then allowed to cool to room temperature. For whole-cell patch clamp recordings, slices were placed on a cover slip coated with poly-L-lysine (Sigma, P4832), transferred to a submersion chamber on an upright microscope (Zeiss AxioExaminer; 40X objective, 1.0 N.A.) and continuously superfused (2-4 ml/min) with warm (∼32-34°C), oxygenated aCSF. Neurons were visualized with a digital camera (Sensicam QE, Cooke) using either infrared differential interference contrast (IR-DIC) microscopy or epifluorescence. Deep L6 neurons were identified using one of the following approaches: 1) neurons retrogradely labeled by stereotaxic injection of either CTB or retrobeads into L1/2; 2) tdTomato-negative neurons with oval shaped cell bodies located below the band of Ntsr1-Cre;tdTomato cells or 3) tdTomato-positive cortical neurons in deep L6 in Nxph4-CreER;tdTomato mice.

### Whole-cell patch clamp recordings

Glass recording electrodes (2-4 MΩ), pulled from borosilicate glass capillaries (1.1 mm ID with filament, 1.5 mm OD, BF150-110-10, Sutter Instruments) on a Sutter P-97 puller, were filled with an internal solution containing (in mM): 2.7 KCl, 120 KMeSO4, 9 HEPES, 0.18 EGTA, 4 MgATP, 0.3 NaGTP, 20 phosphocreatine (Na), pH 7.3, 295 mOsm. In a subset of experiments, the internal solution included 0.25% w/v biocytin (Sigma, B4261) to assess the recorded cell’s laminar location and morphology. Whole-cell patch clamp recordings were obtained using a Multiclamp 700B amplifier (Molecular Devices) and digitized using an ITC-18 (Instrutech) controlled by software written in Igor Pro (Wavemetrics, version 6.1.2.1). All signals were low-pass filtered at 10 kHz and sampled at 20-100 kHz. The series resistance (12.7 ± 5.5 MΩ (mean ± SD), all < 30 MΩ; n=162 cells) was not compensated, and recordings were not corrected for the liquid junction potential. The resting membrane potential was measured after whole-cell configuration was achieved. Neurons exhibiting a resting membrane potential greater than –60 mV were excluded from electrophysiological analyses. The input resistance was determined by measuring the voltage change in response to a 1 s-long –100 pA hyperpolarizing current step. The rheobase was determined using 1 s long depolarizing current steps to evoke single action potentials. Action potential properties were measured from single spikes evoked by the rheobase current injections. Not all measurements were performed for each recorded cell.

### Channelrhodopsin-2 assisted circuit mapping

ChR2 was expressed in geniculocortical neurons as described under *Stereotaxic injections of retrograde tracers and viral constructs*. To express ChR2 in L5 pyramidal neurons, Rbp4-Cre mice (Gerfen et al., 2013) were crossed with a Cre-dependent ChR2-YFP line (Ai32) (Madisen et al., 2012). To express ChR2 in L6CThNs, Ntsr1-Cre mice (Bortone et al., 2014; Guo et al., 2017; Kim et al., 2014) were crossed with the Cre-dependent ChR2-YFP line. Recordings were performed at P26-P45 (GC input: P37-P45; L5 input: P31-P42; L6CThN input: P26-P40). To compare the relative synaptic strength of thalamic or cortical input onto L6a and L6b neurons, two or ten pulses (3 ms) of a small spot of blue light (470 nm, Luminous LED) were delivered to a circular area (∼200-250 μm diameter) containing the pair of recorded neurons to photoactivate ChR2 (Kim et al., 2014; Petreanu et al., 2007). Because the light intensity was adjusted so as not to evoke action potentials (0.5-600 mW/mm^2^), the absolute magnitudes of the responses cannot be compared across experiments. At least 10 traces of synaptic responses were averaged for analysis. All recordings were performed with pairs of neurons to compensate for any differences in ChR2 expression or stimulation level across slices and regions within a slice.

In a subset of the optogenetic experiments, after measuring the synaptic responses of the neuron pairs to ChR2 photostimulation, glutamatergic receptor blockers were bath applied to confirm that the responses were mediated by glutamate. 5 μM 2,3-Dioxo-6-nitro-1,2,3,4-tetrahydrobenzo[f]quinoxaline-7-sulfonamide disodium salt (NBQX, AMPA receptor antagonist, Tocris, 1044) and 5 μM (RS)-3-(2-Carboxypiperazin-4-yl)-propyl-1-phosphonic acid (CPP, NMDA receptor antagonist, Tocris, 0173) were added to the aCSF and perfused for 3 min after which the synaptic responses of the neurons to photostimulation were measured again. The drugs were then washed out by perfusing normal aCSF. Following the recordings, both DIC and fluorescence high (40x, 1.0 N.A.) and low (5x, 0.16 N.A.) magnification images of the recorded cells were taken to confirm their laminar location and cell identity based on the presence or absence of fluorescent markers.

### Morphological reconstruction and analysis

To visualize the morphology of biocytin-filled neurons, brain slices containing the biocytin-filled neurons were fixed with 4% PFA in 0.01 M PBS at 4°C for at least 1 day following electrophysiological recordings. To visualize the neurons, two methods were used: diaminobenzidine (DAB) processing or visualization with fluorescence-conjugated streptavidin. For the DAB method, brain slices were first rinsed in 0.01 M PBS, and endogenous peroxidases were quenched in 1% hydrogen peroxide (H_2_O_2_, Fisher Chemical, H325-500) and 10% methanol in 0.01 M PBS for 30 min. After rinsing in 0.01 M PBS, slices were then permeabilized in 2% Triton X-100 in 0.01 M PBS for 1 hr. Slices were then treated with an avidin-biotin complex solution (ABC, Vectastain, PK-6100) composed of 1% Reagent A with 1% of Reagent B in 2% Triton X-100 in 0.01 M PBS at least overnight at 4°C. After rinsing the sections three times for 10 min in 0.01 M PBS, slices were incubated in 0.05% DAB solution (Sigma, D8001) with 0.033% H_2_O_2_ in 0.01 M PBS for approximately 5 min or until the slice turned light brown.

Slices were then rinsed three times for 10 min in 0.01 M PBS and subsequently darkened via a one min incubation in 0.1% osmium (osmium tetroxide, Electron Microscopy Sciences, 19152) in 0.01 M PBS. Slices were then rinsed three times for 10 min in 0.01 M PBS before being mounted onto glass slides and coverslipped in mowiol mounting media (Mowiol 40-88, Sigma, 324590). For DAB staining, biocytin-filled neurons were visualized on an Imager M2 microscope (Carl Zeiss) using 2.5x (0.075 N.A.), 20x (0.5 N.A.), or 100x (1.4 N.A.) objectives. Three-dimensional reconstruction of filled neurons was performed with Neurolucida (MicroBrightfield) on a Zeiss M2 microscope (100x objective lens, 1.4 N.A.). When using fluorescence-conjugated streptavidin to visualize the biocytin, brain slices were incubated in 2% Triton X-100 in 0.01 M PBS for 1 h at room temperature, and then incubated with AlexaFluor 647-conjugated streptavidin (1:200, S32357, ThermoFisher Scientific) for 24 h at 4°C. Slices were rinsed again and mounted in Vectashield (Vector Laboratories, H-1000-10) or Aqua/Poly Mount mounting medium. Images of processed neurons were obtained on a confocal microscope (LSM 510, Zeiss, 40x objective lens, 1.3 N.A. and/or LSM800, Zeiss, 10x, 0.3 N.A. or 20x, 0.8 N.A.). A subset of recorded neurons was tested for CTGF expression following electrophysiological recordings. After recording, slices were fixed in 4% PFA for at least 1 day and immunostained for CTGF (goat anti-CTGF, 1:300, Santa Cruz Biotechnology, SC-14939; AlexaFluor 488-conjugated Donkey anti-Goat IgG (H+L) antibody, 1:300, A11055, ThermoFisher Scientific) as described above. Biocytin was visualized using AlexaFluor 647-conjugated streptavidin as described above. To define the normalized location of layer boundaries, we used Ntsr1-Cre;tdTomato mice (Somatosensory cortex: n = 3, P37; Visual cortex: n = 4, P82-124). The upper boundary of Cre-expressing somata was designated as the border between L5 and L6, while the upper border of the band of bright Cre-expressing neurites designated the border between L4 and L5a (Kim et al., 2014).

### Single cell RNA-sequencing analysis

Single cell RNA-sequencing (scRNA-seq) datasets for mouse ALM and V1 were obtained from the Allen Institute (Tasic et al., 2018) and downloaded from the NCBI Gene Expression Omnibus (Accession GSE115746). The 10x Genomics single-nuclei RNA-seq (snRNA-seq) dataset for mouse M1 was obtained from the Broad Institute (Yao et al., 2021) and downloaded from the NeMO Archive (RRID:SCR_016152) under “BICCN_Mop_snRNA_10X_v3_Broad.” Cells annotated as low-quality cells or doublets by their respective sources were removed from the datasets. The following gene expression matrices were analyzed: ALM (8,288 cells by 45,768 genes), V1 (13,586 cells by 45,768 genes), and M1 (159,738 cells by 31,053 genes). To visualize the cells, Monocle3 (Cao et al., 2019; Qiu et al., 2017; Trapnell et al., 2014) was used for principal components analysis (PCA) and uniform manifold approximation and projection (UMAP) embedding (McInnes et al., 2018). For the M1 dataset, batch alignment was performed with the mutual nearest neighbors algorithm (Haghverdi et al., 2018). ALM and V1 datasets could not undergo batch alignment due to layer-enriching dissections, labeled cells isolated from transgenic lines, and retrogradely labeled cells that were not evenly distributed across all batches. Cell type annotations correspond to the “cell_subclass” metadata in the ALM and V1 datasets and the “subclass_label” metadata in the M1 dataset.

### Quantification and statistical analysis

All analyses were performed in Igor Pro, MATLAB, Microsoft Excel or ImageJ (Schneider et al., 2012). Data are presented as the mean ± SEM unless otherwise noted. All statistical tests were two-tailed. The Wilcoxon rank sum test and Wilcoxon signed rank test were used to test for statistical significance as noted in the text. To determine the relationship between age and electrophysiological properties, we fit a linear regression model function (StatsLinearRegression; Igor) to perform a linear regression between age and the electrophysiological value across the population of neurons. The same approach was used to determine the relationship between age and CTGF neuron percentage. To compare the short-term synaptic plasticity of L5 and L6CThN input to L6b neurons, we used a two-way ANOVA.

## Acknowledgements

This work was supported by two Brain and Behavior Research Foundation Young Investigator Awards (SPB, JK), R21 MH128765 (JK), a Klingenstein-Simons Fellowship in the Neurosciences (SPB), National Science Foundation (NSF) Grant 1656592 (SPB, LAG), NIH Training Grant 5T32EY017203 (CJM, TAB), two Johns Hopkins Provost’s Undergraduate Awards (ML, EMS), a Kavli Neuroscience Discovery Institute Distinguished Graduate Student Fellowship (ACS), a NSF Predoctoral Fellowship (CJM), a Boehringer-Ingelheim Fonds Fellowship (MC), R01 DC009607 (POK), P30 NS050274, the KBRI Basic Research Program (24-BR-02-02) and a National Research Foundation of Korea grant (NRF-2023-00248148) funded by the Ministry of Science and ICT.

## Author contributions

S.-J.K., J.K. and S.P.B. conceived of the study; S.-J.K., T.A.B., K.L., C.J.M, E.M.S., A.E.C., M.P., H.K., J.A.K., M.C., K.L., and J.K. performed the experiments; S-J.K., T.A.B., A.C.S, M.L., E.M.S., A.E.C, L.A.G., J.K., and S.P.B. analyzed the data. P.O.K. contributed a mouse line. S.-J.K., J.K. and S.P.B. wrote the paper with input from all authors.

